# The immune profile of circulating autoreactive CD4 T cells is imprinted through tissue activation during autoimmune liver diseases

**DOI:** 10.1101/2024.03.26.586770

**Authors:** Anaïs Cardon, Thomas Guinebretière, Chuang Dong, Laurine Gil, Sakina Ado, Pierre-jean Gavlovsky, Martin Braud, Richard Danger, Christoph Schultheiß, Aurélie Doméné, Perrine Paul-Gilloteaux, Caroline Chevalier, Laura Bernier, Jean-Paul Judor, Cynthia Fourgeux, Astrid Imbert, Marion Khaldi, Edouard Bardou-Jacquet, Laure Elkrief, Adrien Lannes, Christine Silvain, Matthieu Schnee, Florence Tanne, Fabienne Vavasseur, Lucas Brusselle, Sophie Brouard, William W Kwok, Jean-François Mosnier, Ansgar Lohse, Jeremie Poschmann, Mascha Binder, Jérôme Gournay, Sophie Conchon, Pierre Milpied, Amédée Renand

## Abstract

Autoimmune liver diseases (AILD) are immune-mediated disorders in which CD4 T cells play a central role. However, the link between circulating self-antigen-specific CD4 T cells and the targeted tissue has not been extensively studied in AILD. We hypothesized that circulating autoreactive CD4 T cells were clonally and functionally related to dominant intra-hepatic pathogenic CD4 T cell clones. Single cell transcriptomic analysis of circulating self-antigen-specific CD4 T cells revealed a specific B-helper and immuno-exhausted transcriptional profile, which was conserved for different autoantigens, but distinct from several other types of foreign antigen specificities. In the blood, the dominant hepatic CD4 T cell clones had a similar transcriptomic signature and were enriched in the PD-1^+^ TIGIT^+^ HLA-DR^+^ CD4 T cell subset. In a mouse model, antigen-specific CD4 T cells acquired the immuno-exhausted transcriptional profile when they accumulated in the liver after local antigen reactivity. Locally, immune checkpoint molecules controlled the response of antigen-specific CD4 T cells responsible for liver damage. Our study reveals the origin and biology of liver-derived autoreactive CD4 T cells in the blood of AILD patients that are imprinted by the liver environment, and suggest a dysregulation of the immune checkpoint molecules pathways. Our study enables tracking and isolating circulating autoreactive CD4 T cells for future diagnostic and therapeutic purposes.

## Introduction

Autoimmune liver diseases (AILD) are rare immune-mediated chronic inflammatory diseases. The three main AILD are autoimmune hepatitis (AIH), primary biliary cholangitis (PBC) and primary sclerosing cholangitis (PSC), which are characterized by the destruction of the liver parenchyma, the intrahepatic biliary epithelial cells of small bile ducts or the intra- or extra-hepatic biliary epithelial cells of the large bile ducts, respectively. AIH, PBC and PSC are characterized by immune infiltrates in the liver^1–7^. Autoantibodies are detectable in the serum of patients, especially for AIH and PBC. PSC is a more complex disease often associated with ulcerative colitis^8^. Although all three AILD are classified as autoimmune diseases, the degree of intensity of the autoreactive response appears to be variable, and the link with the autoimmune T-cell signature is unknown. In our study, we focused primarily on AIH and PBC diseases.

In AILD, the strong association with the HLA locus, the presence of autoantibodies in serum and T cells in damaged tissue reflects the complete adaptive immune response against self-antigens, with a central role for CD4 T cells with their helper function^9–12^. The recent demonstration of the efficacy of rituximab (anti-CD20) treatment in AIH confirms that B cells contribute to the pathogenesis of AILD^13^. However, it has been difficult to establish in-depth the immune profile of autoreactive CD4 T cells, as they are rare in the blood and not all self-antigens are known. During AILD, some target self-antigens have been described. Anti-SLA antibodies target the Sepsecs self-antigen, anti-LKM1 target the CYP2D6 self-antigen, anti-LC1 target the FTCD self-antigen and the anti-mitochondrial M2 (anti-M2) antibody target the PDCE2 self-antigen. T cell reactivity against Sepsecs, CYP2D6, FTCD and PDCE2 is detectable in the blood of AILD patients with a preferential Th1 cytokine profile (IFNγ)^14–17^. In a mouse model, immunization against CYP2D6 or FTCD induces liver autoimmunity^18–21^. Those studies reinforce the idea of a complete adaptive immune response against self-antigens in AILD with a central role of CD4 T cells. Recently, we were able to characterize the single cell transcriptomic profile of Sepsecs-specific CD4 T cells in the blood of AIH patients with anti-SLA antibodies^22^. Compared to Candida Albicans-specific CD4 T cells, Sepsecs-specific CD4 T cells were PD-1^+^ CXCR5^-^ cells and expressed high level of the B-helper gene, *IL21*, as well as the *IFNG* gene and the immunoregulatory genes *TIGIT* and *CTLA-4,* among others. In the blood of AIH patients, we found an increase in IL21- and IFNγ-producing PD-1^+^ CXCR5^-^ CD4 T cells, independent of the presence of specific autoantibodies (e.g anti-SLA)^22^. The signature of these autoreactive CD4 T cells in the blood presented high similarity with the peripheral T helper cells (T_PH_) identified in the synovium of patients with rheumatoid arthritis and in other tissues during autoimmunity^23–29^. However, it was still unknown whether the T_PH_ immune signature was a common feature of other liver-self-antigen-specific CD4 T cells.

The similarity between circulating Sepsecs-specific CD4 T cells and T_PH_ in the tissue raised the question whether circulating autoreactive CD4 T cells are directly linked to the reactivity in the tissue. Although T_PH_ cells can also be found in smaller numbers in the blood, the site of their differentiation is unknown^22,23,30^. In liver biopsies from AIH and PBC patients, Sepsecs- and PDCE2-reactive T cells are detectable but poorly characterized; they produce TNF and/or INFγ^31–34^. Characterizing the immune signature of T cells in tissues is a timely topic, but the dynamic link between tissues and circulation is less well understood, particularly for CD4 T cells. Liver biopsies are small and rare, making it difficult to characterize liver-autoreactive CD4 T cells during AILD in the tissue. Thus, the capacity to detect in the blood autoreactive cells directly linked to tissue pathogenesis is an interesting perspective. Following an immune response or infection, peripheral CD4 and CD8 T cells can migrate into tissues to differentiate into tissue-resident memory (T_RM_) cells that are characterized by clonal retention and segregation in the tissue^35,36^. Interestingly, T_RM_ CD4 T cells, T_PH_ cells and circulating Sepsecs-specific CD4 T cells express high levels of immune checkpoint molecules (PD-1, TIGIT and CTLA-4). This may suggest a tissue imprint of T_PH_ and Sepsecs-specific CD4 T cells found in the peripheral circulation due to chronic antigen stimulation in the tissue. We thus hypothesized that some circulating autoreactive CD4 T cell derive from an exacerbated clonal expansion in the tissue during active autoimmunity.

Here we used integrative single-cell immuno-transcriptomics to characterize three distinct liver-self-antigen-specific CD4 T cell reactivities and expanded intrahepatic CD4 T cell clonotypes in the blood of AILD patients. We revealed a shared signature and demonstrated that liver-autoreactive CD4 T cells can be detected in the peripheral bloodstream. In a mouse model designed to study the initiation of an immune response against an antigen expressed in the liver, we observed alterations of the transcriptomic profile of intrahepatic antigen-specific CD4 T cells after local antigen reactivity. The most relevant markers characterizing both intrahepatic and circulating liver-autoreactive CD4 T cells were the immune checkpoint molecules TIGIT, PD-1 and CTLA-4, which were involved in the control of local antigen-specific T cell reactivity. These results demonstrate how circulating liver-self-antigen-specific CD4 T cells are linked to tissue auto-reactivity and how the hepatic environment imprints their immune signature, with an important clinical perspective for monitoring and targeting liver auto-reactivity.

## Results

### Self-antigen-specific CD4 T cell detection in AILD patients with auto-antibodies

We have previously shown that Sepsecs (SLA)-specific CD4 T cells are detectable in the blood of patients with autoimmune hepatitis (AIH) whose serum contains anti-SLA antibodies^22^. In the present study, we investigated whether we could extend those findings to autoreactive CD4 T cells specific for other antigens frequently targeted by autoantibodies in AILD. We studied the reactivity of CD4 T cells against Sepsecs, CYP2D6 and PDCE2 in AILD (AIH, PBC and overlap AIH/PBC) patients with or without anti-SLA (SLA^+^) or anti-LKM1 (LKM1^+^) or anti-M2 (M2^+^) autoantibodies. As an experimental control, we also studied the reactivity of CD4 T cells against MP65 (Candida Albicans; C.ALB), MP1 (Influenza; H1N1) and SPIKE (SARS-CoV-2) in the AILD patients. As previously described^22^, detection of antigen-specific CD4 T cells was achieved after four hours peptide stimulation *in vitro*. CD4 T cells that specifically recognize antigen-derived peptides upregulate CD40 ligand (CD154) and can be detected by flow cytometry (Figure 1A and B). These reactive CD4 T cells were memory cells (CD45RA^-^; mCD4 T cells) expressing high levels of PD-1 but not CXCR5 (Figure 1B and C). The frequency of Sepsecs-, CYP2D6- and PDCE2-specific PD-1^+^ CXCR5^-^ mCD4 T cells was significantly higher in patients with the presence of specific autoantibodies (Figure 1D). In comparison, the frequency of MP1 (H1N1)-, MP65 (C.ALB)- and SPIKE (SARS-CoV-2)-specific CD4 T cells was detectable in the blood of all patients (Figure 1E). Foreign antigen-specific CD4 T cells were also mCD4 T cells. MP1- and SPIKE-specific CD4 T cells expressed PD-1. Neither MP1-, MP65- or SPIKE-specific CD4 T cells expressed significant levels of CXCR5 (Figure 1E). In conclusion, during AILD, we observed an association between the specific autoimmune humoral response and the presence of circulating self-antigen specific CD4 T cells.

**Figure 1.**
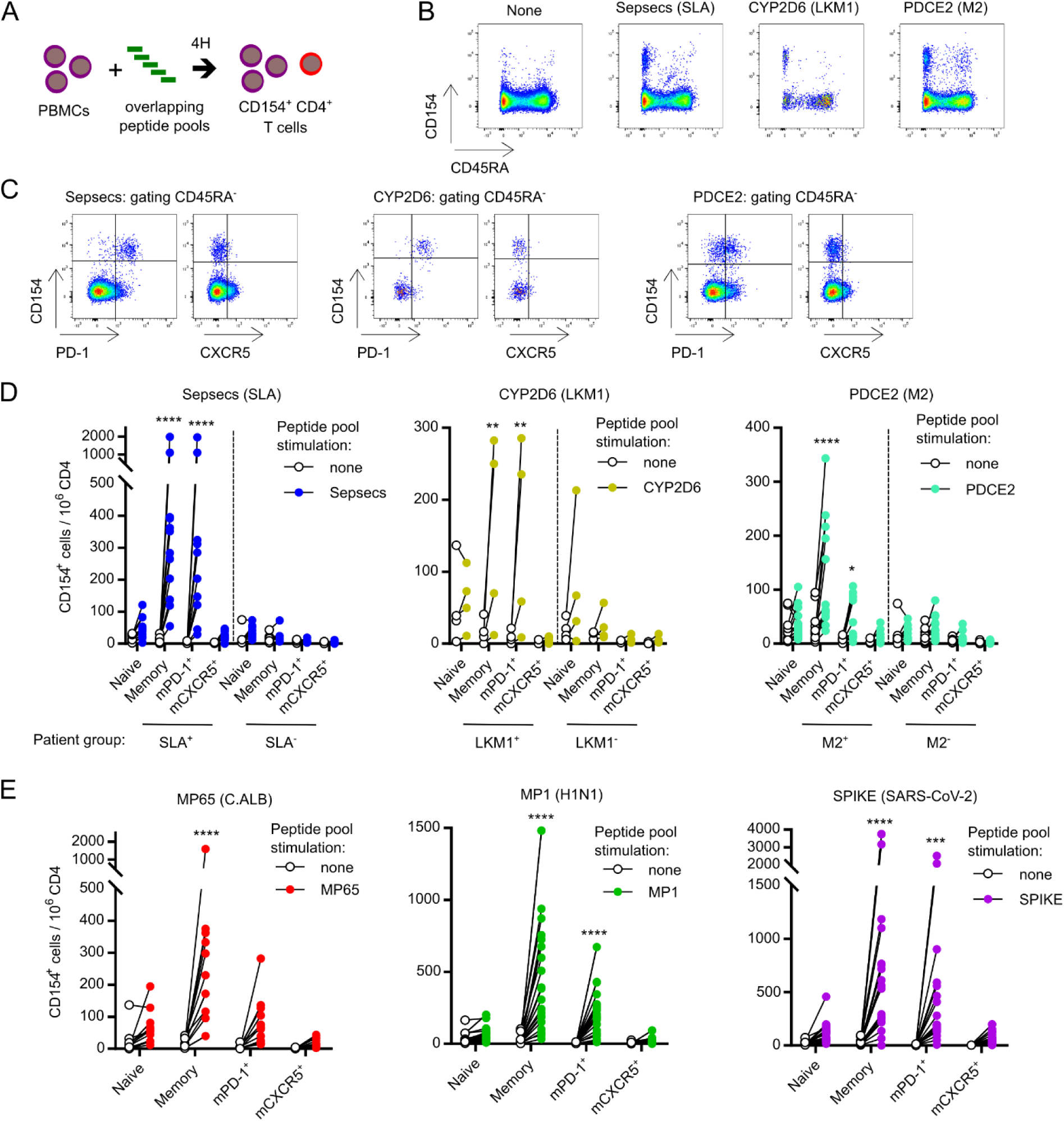
Detection and immune-phenotyping of circulating liver-self-antigen-specific CD4 T cells. (A) Schematic representation of the detection of antigen-specific CD4 T cells based on the upregulation of CD40 ligand (CD154) after 4 hours of *ex vivo* peptide stimulation. (B) Pseudocolor dot plot representation of CD45RA and CD154 expression at the surface of CD4 T cells after stimulation with peptide pools from the indicated antigens. (C) Pseudocolor dot plot representation of PD-1, CXCR5 and CD154 expression at the surface of CD45RA^-^ mCD4 T cells after stimulation with peptide pools from the indicated antigens. (D) Frequency of CD154^+^ naïve, memory, memory PD-1^+^ (mPD-1^+^) and memory CXCR5^+^ (mCXCR5^+^) CD4 T cells per million CD4 T cells after Sepsecs, CYP2D6 or PDCE2 peptides stimulation of PBMCs from 12 SLA^+^ patients and 9 SLA^-^ patients (left); or 4 LKM1^+^ patients and 4 LKM1^-^ patients (central); or 14 M2^+^ patients and 14 M2^-^ patients (right). (E) Frequency of CD154^+^ naïve, memory, mPD-1^+^ and mCXCR5^+^ CD4 T cell per million CD4 T cells after MP65 (C.ALB), MP1 (H1N1) or SPIKE (SARS-CoV-2) peptides stimulation of PBMCs from 13, 24 and 23 patients respectively. Sidak’s multiple comparisons test was used for D and E. *: p<0.05; **: p<0.01; ***: p<0.001; ****: p<0.0001.

### Self-antigens-specific CD4 T cells have a unique transcriptomic immune signature during AILD

We sorted self-antigen-specific and foreign antigen-specific mCD4 T cells by flow cytometry for single-cell RNA-seq and TCR-seq analyses (Figure 2). An unsupervised clustering analysis led to the identification of eight clusters (Figure 2A). The majority of transcriptional clusters corresponded to distinct antigenic specificities rather than to distinct patients (Figure 2B and C). The only notable exception was cluster 7, which corresponded to a PDCE2-specific CD4 T cell clonotype from one patient (01-176) with a unique cytotoxic profile (Supplementary Figure 1).

**Figure 2.**
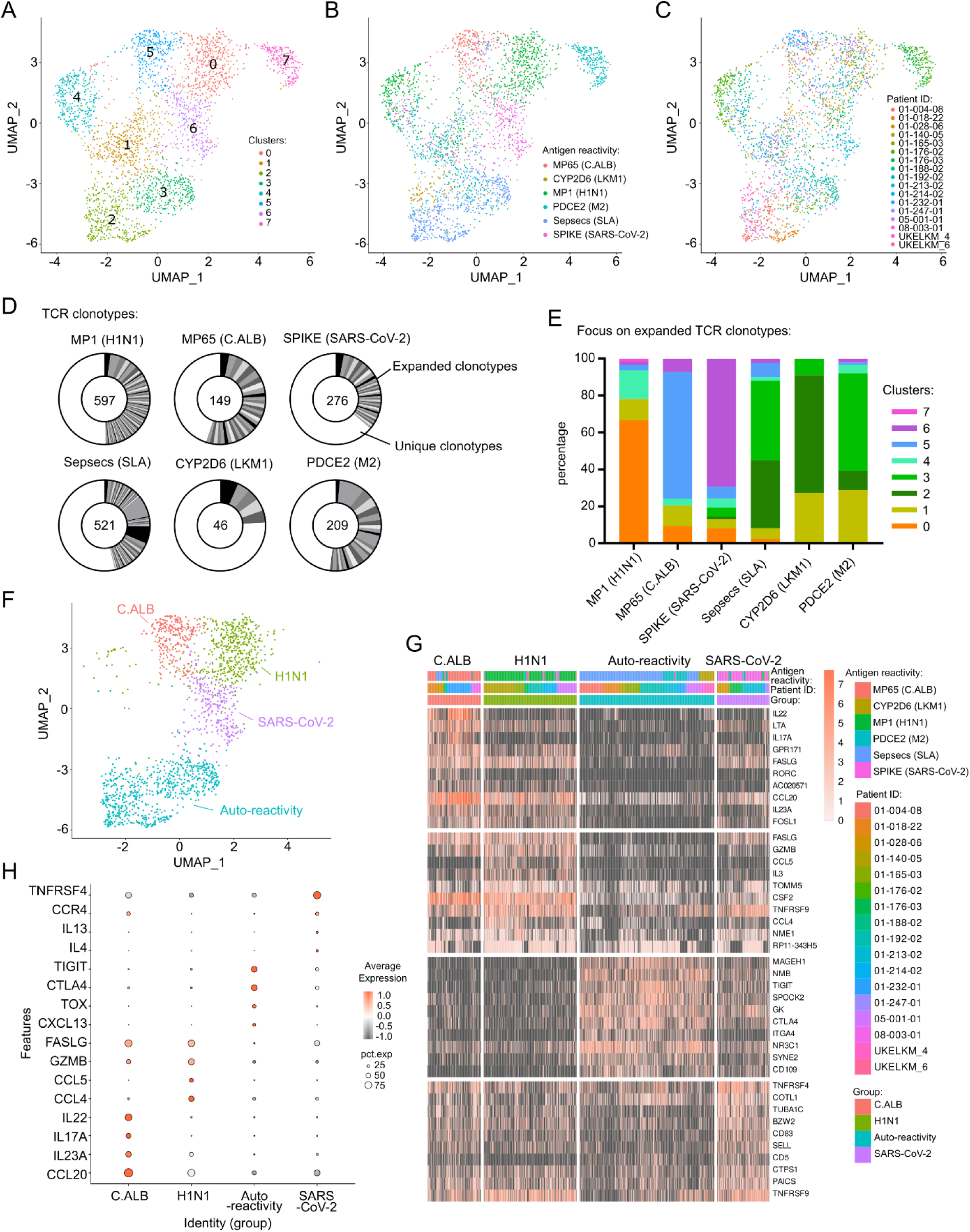
Single-cell RNA sequencing of liver-self-antigens specific CD4 T cells. A total of 2768 CD154^+^ mCD4 T cells (648 Sepsecs-, 113 CYP2D6-, 583 PDCE2-, 273 MP65-, 798 MP1- and 353 SPIKE-specific CD4 T cells) from 7 SLA^+^, 3 LKM1^+^ and 6 M2^+^ patients were analyzed for transcriptome and TCR sequences at the single cell level using FB5P-seq. (A) UMAP representation of antigen-specific memory CD4 T cell single-cell transcriptomes, colored by non-supervised Louvain cluster identity. (B) UMAP representation colored by antigen reactivity. (C) UMAP representation colored by patient ID. (D) TCRαβ clonal diversity of antigen-specific single T cells for each antigen-reactivity. Numbers indicate the number of single cells analyzed with a TCRαβ sequence. Black and grey sectors indicate the proportion of TCRαβ clones (clonotype common to ≥ 2 cells) within single-cells analyzed; white sector: unique clonotypes. (E) Louvain cluster distribution of antigen-specific T cell clonotypes for each reactivity, as indicated. (F) UMAP representation of selected antigen reactivity cluster groups of memory CD4 T cells (C.ALB, H1N1, SARS-CoV-2 and auto-reactivity). (G) Single-cell gene expression heatmap for top 10 marker genes of cells from each antigen reactivity group defined in (F). (H) Dot plot representation of four selected marker genes of clusters C.ALB, H1N1, SARS-CoV-2 and auto-reactivity.

As the memory response against an antigen is generally associated with clonal expansion, we analyzed the diversity and proportion of TCR clonotypes for each antigen reactivity (Figure 2D). Clonal expansion was observed for every antigenic reactivity, whether associated with self- or foreign-reactivity. Analysis of these expanded clones for cluster affiliation demonstrated that transcriptional clusters were linked to antigenic reactivity (Figure 2E). Interestingly, all three liver auto-reactivities were grouped in the same clusters. Sepsecs-, CYP2D6- and PDCE2-specific CD4 T cells were in the clusters 2 and 3. MP1-specific CD4 T cells were in the cluster 0; MP65-specific CD4 T cells were in the cluster 5; and SPIKE-specific CD4 T cells were in the cluster 6. Two clusters (4 and 1) were not linked to antigenic reactivity. We annotated four antigen-reactivity groups based on unsupervised transcriptional clustering: the H1N1 cluster (ex-cluster 0); the C.ALB cluster, (ex-cluster 5); the SARS-CoV-2 cluster (ex-cluster 6); and the auto-reactivity cluster (ex-clusters 2 and 3) (Figure 2F and G). The gene signature of autoantigen-reactive cells was linked to the regulation of T cell activation and immune exhaustion (*CTLA-4, TIGIT, BATF, IL21, CD74, PRDM1, TOX*). This gene signature, together with the *CXCL13* gene, was also linked to the B-helper autoreactive signature of T_PH_ described in other autoimmune disorders (Figure 2H). In comparison, the H1N1 gene signature was associated with a classical antiviral response and T cell chemotaxis (*GZMB*, *CCL4, CCL5 and FASLG*). The Candida Albicans gene signature was associated with a type 17 helper response (*CCL20, IL22, IL17A, IL23A*). SARS-CoV-2 gene signature was associated with a type 2 helper response and B cell activation (*TNFRSF4, CCR4, IL4 and IL13*). We had the opportunity to longitudinally analyze the TCR clonotypes of Sepsecs-specific and MP65-specific CD4 T cells of patients (n=3 and 1, respectively) over 3 years (2018 to 2021). Although variable numbers of novel clones were detected in patients’ blood over time, some conserved Sepsecs-specific CD4 T clonotypes were detected over the 3-year period (Supplementary Figure 2), suggesting chronically active clonal expansion of autoreactive CD4 T cells despite their immuno-exhausted molecular profile. Altogether, these data demonstrate that distinct liver-self-antigen-specific CD4 T cells had a common specific B-helper autoreactive signature, with high-level expression of checkpoint inhibitors (immuno-exhausted transcriptional profile).

### Clonal overlap between circulating autoreactive and intra-hepatic CD4 T cells in AILD

We then attempted to answer the following question: how related to liver infiltrating CD4 T cells are circulating autoreactive CD4 T cells in AILD? We hypothesized that circulating autoreactive CD4 T cells might represent clonal relatives of liver CD4 T cells and that their specific molecular profile was induced during chronic antigen reactivity in the tissue. For one SLA^+^ patient analyzed by peptide restimulation and single-cell sequencing (Figure 2, patient 01-192), we also had access to a frozen liver biopsy, from which we derived bulk TCR-beta-seq data (Supplementary Figure 3A). Six out of sixty (10%) Sepsecs-specific circulating CD4 TCR clonotypes were also detected in the liver, compared to only one out of forty-six (2.2%) MP1-specific TCR clonotypes (H1N1). Bulk TCR-beta sequence counts corresponding to Sepsecs-specific clonotypes suggested few clonal expansions dominating the Sepsec-specific response also in the liver (Supplementary Figure 3C). Thus, for this patient, circulating self-antigen-specific CD4 T cells reflected clonally expanding autoantigen-specific T cells within the liver.

For most AILD patients, no self-antigen reactive TCR were known. Therefore, to continue exploring the relationship between circulating and liver CD4 T cells, we took advantage of our initial observation that in AILD, most circulating self-antigen-specific CD4 T cells have a PD-1^+^ CXCR5^-^ CD45RA^-^ memory phenotype^22^ (Figure 1). We had access to paired blood-liver samples from 4 AILD patients; for each, we extracted genomic DNA and performed bulk TCR-beta-seq from blood PD-1^+^ CXCR5^-^ mCD4 T cells, blood PD-1^-^ mCD4 T cells, and whole liver biopsy (Supplementary Figure 4A). First, we observed that the 100 most abundant TCR-beta sequences in the liver account for more than 50% of intrahepatic TCRs, suggesting strong local clonal expansion (Supplementary Figure 4B). We also observed that the PD-1^+^ CXCR5^-^ mCD4 T cell population shared more TCR-beta sequences with the liver than PD-1^-^ mCD4 T cells (Supplementary Figure 4C). Interestingly, the proportion of cells with the top 100 liver TCR-beta sequences was high in the PD-1^+^ CXCR5^-^ mCD4 T cell population, than in the PD-1^-^ mCD4 T cells (supplementary Figure 4D). This data demonstrated the circulating PD-1^+^ CXCR5^-^ mCD4 T cell population contain dominant intrahepatic CD4 T cell clonotypes.

Based on this, we performed droplet-based single-cell RNA-seq combined with TCR sequencing of blood PD-1^+^ CXCR5^-^ mCD4 T cells from the four patients to characterize the transcriptional profile of cells with TCR clonotypes detected in the matched liver biopsies (Figure 3A). First, cluster analysis of PD-1^+^ CXCR5^-^ mCD4 T cells revealed the heterogeneity of these cells (Figure 3B). The main subsets identified by unsupervised azimuth annotation and marker gene analysis were: *GZMK*^+^ effector CD4 T cells (T_EM_ GZMK^+^, cluster 1); *HLA-DR*^+^ recently activated central memory CD4 T cell (T_CM_ HLA-DR^+^, cluster 2), *NKG7*^+^ effector CD4 T cell (T_EM_ NKG7^+^, clusters 3); cytotoxic (*GZMB*^+^*GNLY*^+^) CD4 T cells (CTL, clusters 5) and *FOXP3*^+^ (regulatory) CD4 T cell (T_REG_, cluster 6) (Figure 3B and C, and supplementary Figure 5 and 6). Clusters 4, 7 and 8 were likely subsets of T_CM_ but were difficult to further annotate based on marker genes.

**Figure 3.**
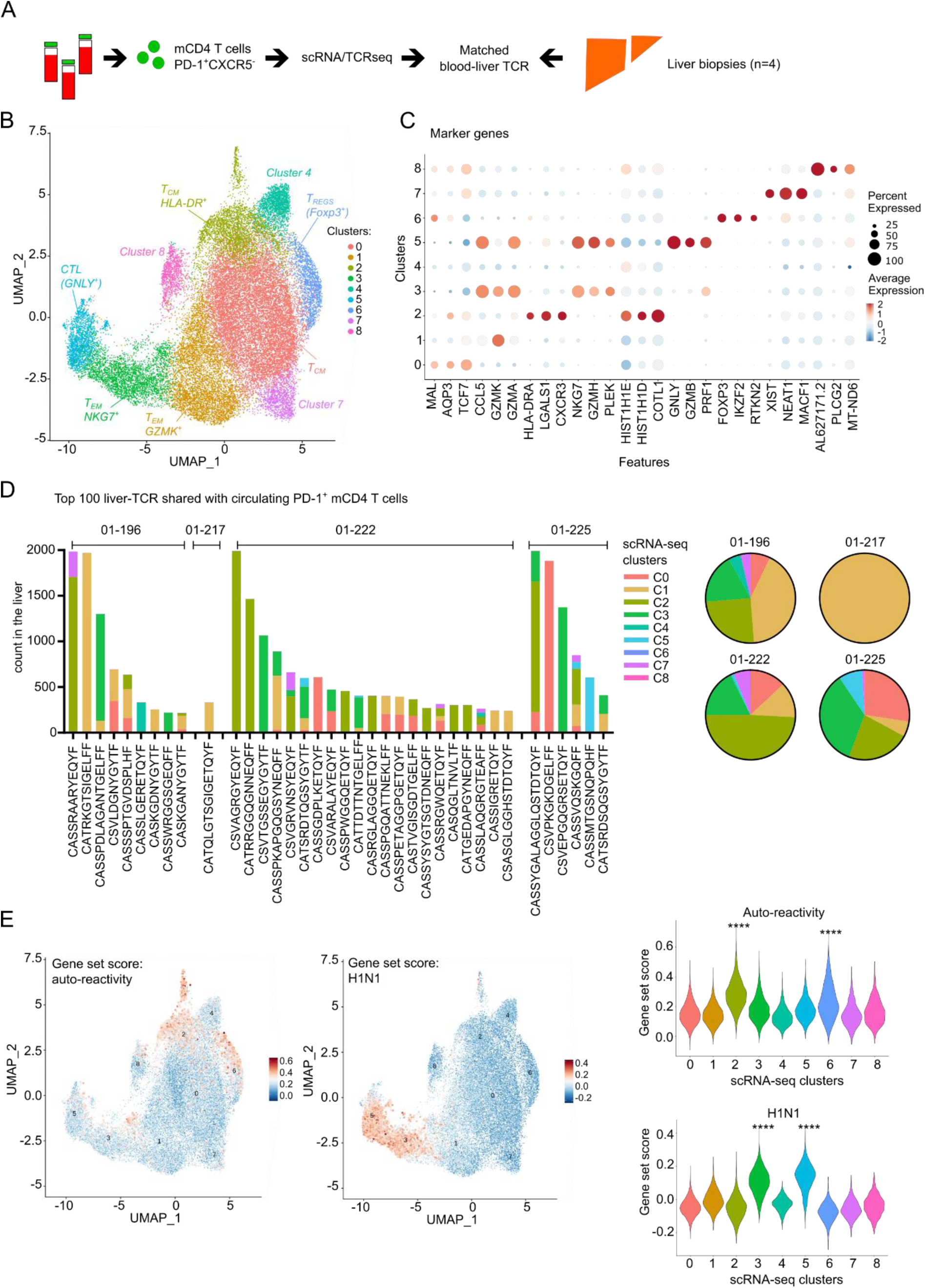
Tracking TCR clonotypes between liver biopsies and circulating CD4 T cell subsets. (A) Experimental design for scRNA-seq and TCR-seq of memory PD-1^+^ CXCR5^-^ CD4 T cells from four distinct AILD patients. (B) UMAP representation of circulating memory PD-1^+^ CXCR5^-^ CD4 T cell transcriptomes, colored by non-supervised Louvain clustering. (C) Dot plot representation of top 3 marker genes of each clusters in (B). (D) Frequency representation (’counts’) of each top 100 largest liver TCRβ sequences found in circulating PD-1^+^ mCD4 T cells. For each TCRβ sequences the cluster affiliation is indicated based on scRNA-seq and TCR-seq of memory PD-1^+^ CXCR5^-^ CD4 T cells. Pie chart represent the global cluster affiliation of the top 100 largest liver TCRβ sequences found in circulating PD-1^+^ mCD4 T cells per patient. (E) Left: UMAP representation of circulating memory PD-1^+^ CXCR5^-^ CD4 T cell transcriptomes colored by gene set score for the indicated antigen-reactivity module. Right: violin plots of gene set score distribution of cells from different Louvain clusters, as indicated. Pairwise comparison with Benjamini-Hochberg corrected Wilcoxon test. ****: p<0.0001.

In this data set, we tracked the top 100 liver TCR-Beta sequences (Figure 3D). For each of these liver TCR-beta clones, we assigned a cluster identification based on single-cell transcriptomic data from PD-1^+^ CXCR5^-^ mCD4 T cells (Figure 3D). The analysis revealed top expanded intrahepatic clones found in the blood were mainly distributed between cluster 1 (T_EM_ GZMK^+^), cluster 2 (T_CM_ HLA-DR^+^) and cluster 3 (T_EM_ NKG7^+^). We computed gene set scores based on our analyses of peptide-stimulated T cells (Figure 2), with the expression of the top 50 genes (avg_log2FC) from auto-reactivity or H1N1 reactivity, respectively. These scores were then quantified in the droplet-based scRNA-seq dataset of non-stimulated PD-1^+^ CXCR5^-^ mCD4 T cells (Figure 3E). Cells in cluster 2 (T_CM_ HLA-DR^+^) had the highest score for the auto-reactivity module, while cells in clusters 3 and 5 had the highest score for the H1N1 module. These data suggest that cells in cluster 2, PD-1^+^ CXCR5^-^ HLA-DR^+^ T_CM_, were the main circulating CD4 T cell clonal counterpart of liver-autoreactive CD4 T cells. Interestingly, this cluster 2 also contained proliferative CD4 T cells, reinforcing the idea of local clonal expansion (Supplementary Figure 6).

We further sought to relate the transcriptional signatures of peptide-stimulated and untouched circulating autoreactive mCD4 T cells (Figure 2 and 3). Based on previously published results^22^, we synthesized HLA-DRB1 03:01 biotinylated monomers loaded with the Sepsecs187-197 epitope peptide, assembled fluorescent-labeled pMHCII tetramers, and monitored Sepsecs-specific CD4 T cells in the blood of six SLA^+^ HLA-DR3^+^ patients (Figure 4A). Peptide-MHCII tetramer positive (TTpos) CD4 T cells were detectable and displayed a PD-1^+^ memory phenotype (Figure 4B). We performed plate-based single-cell transcriptomic analyses of pMHCII TTpos Sepsecs-specific CD4 T cells. We compared pMHCII TTpos and TTneg CD4 T cells for their expression of gene modules associated with antigen reactivities defined in Figure 2 (Figure 4C) and in the PD-1^+^ CXCR5^-^ mCD4 T cell subsets identified in Figure 3 (Figure 4D). As expected, pMHCII TTpos CD4 T cells expressed significantly higher levels of gene modules associated to self-antigen specific CD4 T cells (Figure 4C) and cluster 2 T_CM_ HLA-DR^+^ CD4^+^ cells (Figure 4D).

**Figure 4.**
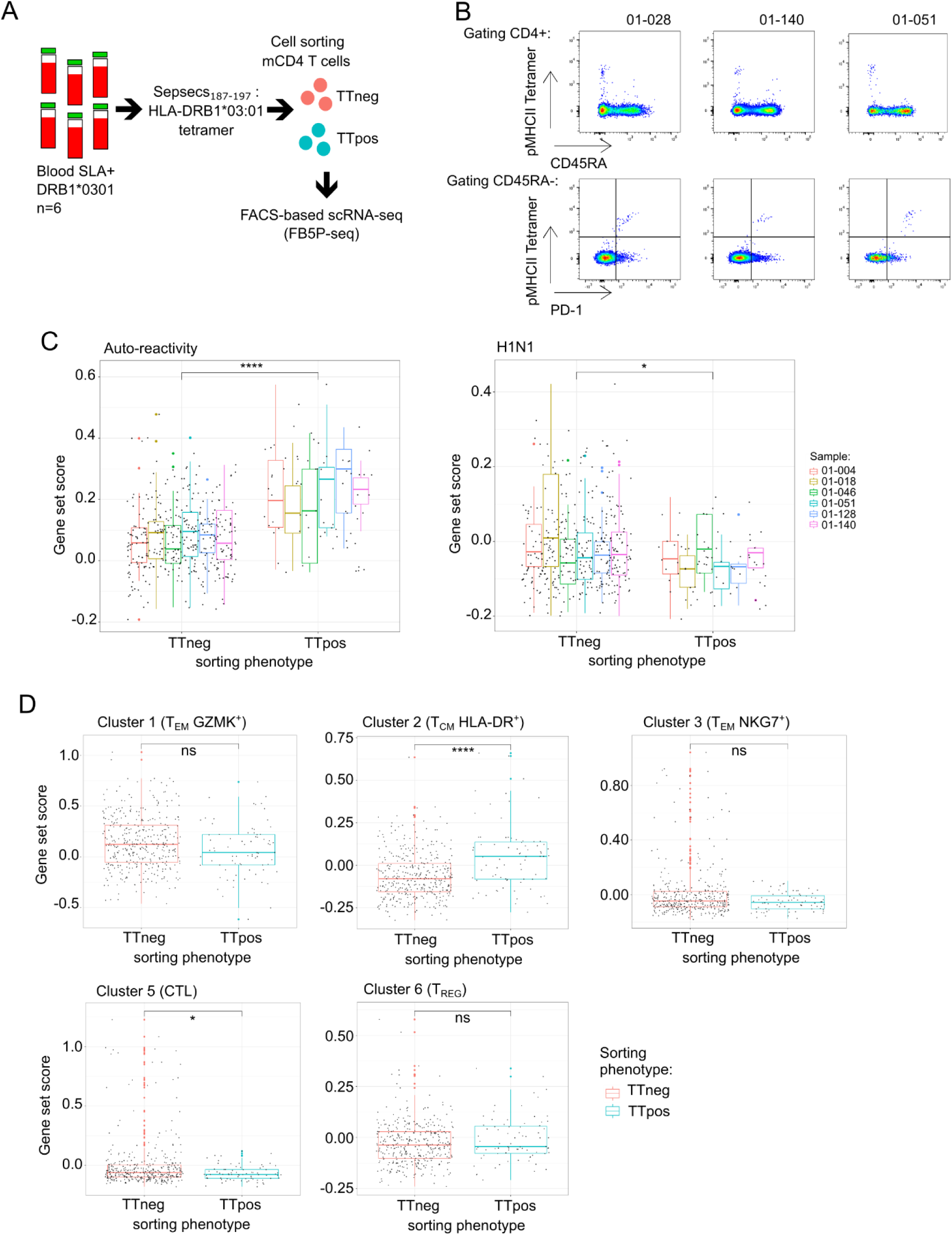
Single cell transcriptomic analysis of unstimulated Sepsecs-specific CD4 T cells. (A) Experimental design for scRNA-seq data analysis of Sepsecs/SLA185-197 HLA-DRB1*0301 pMHCII tetramer positive cells. (B) Pseudocolor dot plot representation of pMHCII tetramer staining in three of six patients. Top: surface expression of CD45RA and pMHCII tetramer in CD4^+^ T cells. Bottom: surface expression of PD-1 and pMHCII tetramer in CD4^+^ CD45RA^-^ T cells. (C) Box plots of gene set score distribution of pMHCII tetramer negative (TTneg) or positive (TTpos) cells for antigen reactivity modules as indicated. (D) Box plots of gene set score distribution of pMHCII tetramer negative (TTneg) or positive (TTpos) cells for top marker genes of circulating memory PD-1^+^ CXCR5^-^ CD4 T cell clusters defined in Figure 3E. Unpaired Mann-Whitney test was used: *: p<0.05; ****: p<0.0001.

Altogether, our integrative analyses of circulating self-antigen-specific and dominant intrahepatic CD4 T cell clones in AILD patients converge to establish that some autoreactive CD4 T cell clones can be found both in the liver and in the blood, where they are defined as PD-1^+^ CXCR5^-^ HLA-DR^+^ mCD4 T cells, with a specific B-helper and immuno-exhausted transcriptional profile.

### High-dimensional phenotyping of circulating liver-autoreactive CD4 T cells

We then examined the precise phenotype of these circulating liver-autoreactive CD4 T cells by spectral flow cytometry using PBMCs from patients affected by different liver diseases: five control non-autoimmune patients with non-alcoholic steatohepatitis (NASH), fourteen AIH patients with an active disease, and thirteen AIH patients in remission under treatment (Supplementary Table 2). We focused on the mCD4 T cell population and performed clustering analysis (Figure 5A-D). The comparison revealed the significant increase of three cell clusters in patients with active AIH in comparison to NASH patients (C6, C12 and C15, Figure 5B). The C6 and C15 clusters were T_CM_ CD27^+^ TIGIT^+^ PD-1^+^ CXCR5^-^ CD49d^+^ CD25^-^ CD127^-^ ICOS^+^ CD4 cells and expressed CD38, a marker we have already observed upregulated by CD4 T cells during active AIH^22^ (Figure 5C and D, Supplementary Figure 7).The C15 cluster was characterized by high HLA-DR expression and may represent the liver-autoreactive CD4 T cells identified previously (Figure 3). This cluster (C15) was high in the blood of active AIH patients compared to AIH patients in remission under treatment (Supplementary Figure 7). The C12 cluster was characterized by CD57 expression but its frequency was very low. We performed a supervised analysis of the HLA-DR^+^ subset and confirmed that it was significantly more frequent during the active phase (AIHa) than in patients with NASH or in remission (AIHr) (Supplementary Figure 8). In parallel, in four active AIH patients, we performed the intracellular analysis of the PD-1^+^ TIGIT^+^ CXCR5^-^ CD4 subset with a new panel of antibodies. We confirmed its heterogeneity; this subset consisted of classical T_REG_ (FOXP3^+^ CD127^-^ EOMES^-^), cytotoxic cells (GZMB^+^ HLA-DR^-^ EOMES^+^ GZMA^+^) and HLA-DR^+^ FOXP3^-^ EOMES^-^ cells (Supplementary Figure 9). This was in agreement with the transcriptomic data (Figure 3) to demonstrate that the liver-autoreactive CD4 T cell cluster (HLA-DR^+^ PD-1^+^) was different from the T_REG_ and cytotoxic (CTL) PD-1^+^ CD4 T cell clusters. Thus, after exclusion of T_REGs_ (using CD127 and CD25 or FOXP3 expression), HLA-DR expression emerged as the most relevant marker for delineating liver-autoreactive CD4 T cells among PD-1^+^ TIGIT^+^ CD4 T cells.

**Figure 5:**
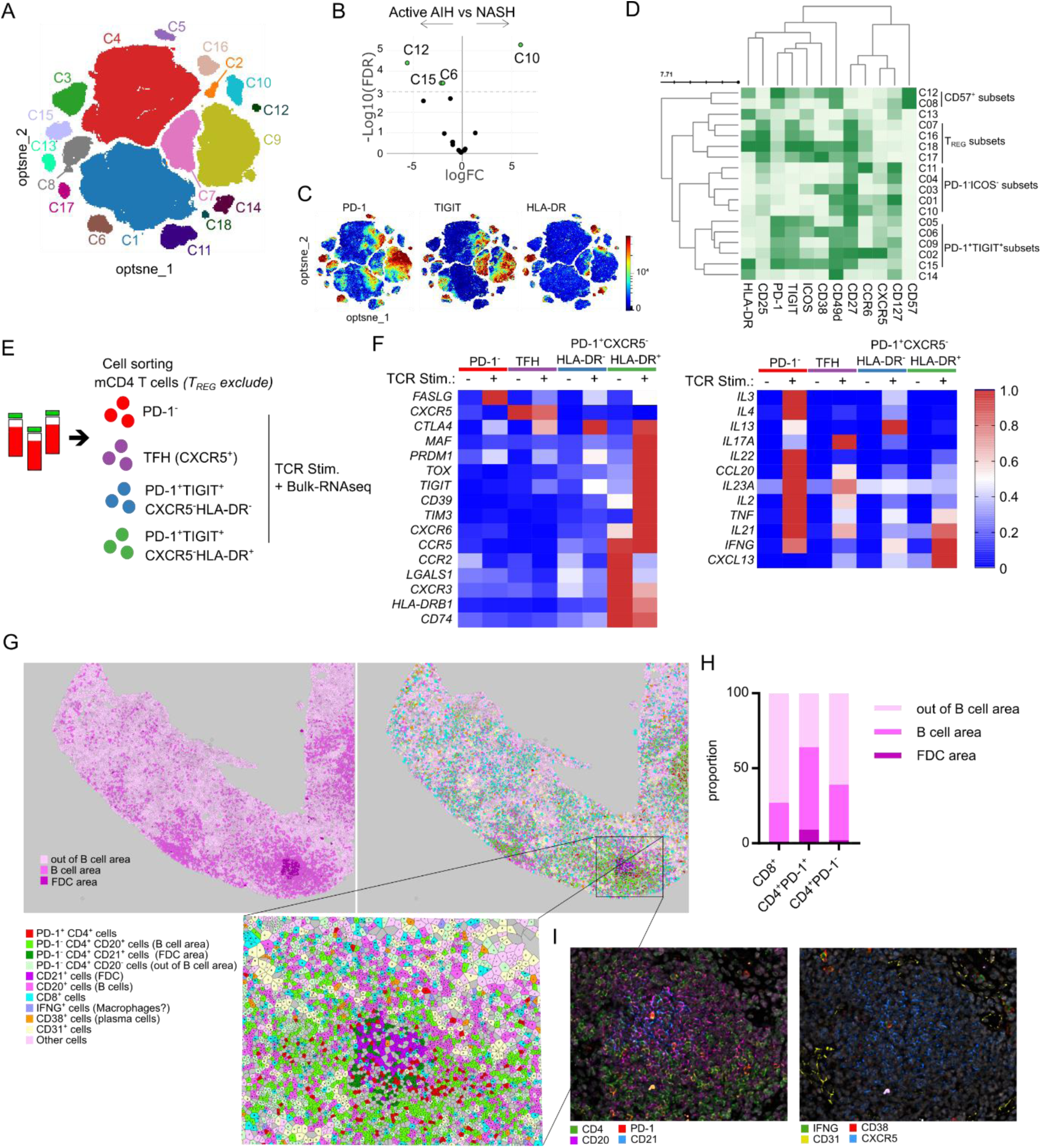
Characterization of circulating PD-1^+^ TGIT^+^ HLA-DR^+^ non-TREG CD4 T cells as liver-autoreactive CD4 T cells. (A) opt-SNE representation of blood memory CD4 T cell subsets, colored by non-supervised FlowSOM clustering, from 5 control non-autoimmune patients with non-alcoholic steatohepatitis (NASH), 14 AIH patients with an active disease (Active AIH), and 13 AIH patients in remission under treatment. (B) Differential cluster abundance between active AIH and NASH patients. (C) PD-1, TIGIT and HLA-DR expression. (D) Surface marker heatmap of the clusters identified in (A). (E) Experimental design for bulk-RNA-seq after cell sorting and 24h *in vitro* TCR stimulation. (F) Selected genes heatmap of indicated CD4 T cell subsets after or not TCR stimulation from five distinct patients (mean representation). (G) Voronoi representation of cellular subsets in an AIH liver biopsy after cell segmentation. FDC area is defined by expression of CD21; B cell area is defined by CD20 expression; and out of B cell area is defined by the absence of CD20 expression. The lower panel shows a zoom on the selection indicated in the upper panel. (H) Representation of the repartition of CD8^+^, CD4^+^ PD-1^+^ and CD4^+^ PD-1^-^ cells in FDC, B cell and out of B cell area defined in (G). (I) CD4, CD20, PD-1, CD21, IFNγ, CD38, CD31 and CXCR5 marker staining within segmentation.

To get an insight into the functionality of these rare cells, we used this strategy of T_REG_ exclusion to sort HLA-DR^+^ PD-1^+^ TIGIT^+^ CXCR5^-^ mCD4 T cells and to perform their TCR stimulation *in vitro* for bulk-RNA sequencing (Figure 5E and F). Like autoreactive CD4 T cells after peptide stimulation, HLA-DR^+^ PD-1^+^ TIGIT^+^ CXCR5^-^ mCD4 T cells significantly upregulated *CXCL13*, *IL21*, *CTLA-4*, *PRDM1*, *TIGIT*, *MAF* and *TOX,* upon TCR stimulation. These cells also expressed *LGALS1*, *CXCR3, HLA-DR and CD74* like CD4 T_CM_ HLA-DR^+^ cells; these genes were downregulated by TCR stimulation. They expressed genes associated to the T_PH_ and T_RM_ CD4 cell signature like *ENTPD1, TIM3, CXCR6, CCR5* and *CCR2* that linked them to the tissue immune response. They could also express *TNF* and *INFG*, but not *IL2*, after TCR stimulation. This subset presented high proximity with autoreactive CD4 T cells and circulating dominant intrahepatic clonotype, thus, we can conclude that HLA-DR^+^ PD-1^+^ TIGIT^+^ CXCR5^-^ CD4 T cells contained the liver-autoreactive CD4 T cells.

We performed a spatial multi-phenotyping analysis on a liver biopsy from one AIH patient, and we identified tertiary lymphoid structures characterized by the presence of B-cells (CD20^+^ cells) and follicular dendritic cells (FDC, CD21^+^) (Figure 5G, and Supplementary Figure 10). Interestingly, PD-1^+^ CD4 T cells were enriched in the FDC and B cell zones, in comparison to the CD8^+^ and the PD-1^-^ CD4 T cells (Figure 5H and I, Supplementary Figure 10). The FDC area was enriched in CXCR5^+^ cells suggesting germinal center formation and/or B cell retention (Figure 5I). Although the phenotype of the PD-1^+^ CD4^+^ T cells was less detailed, this localization suggested that liver-autoreactive CD4 T cells could be involved in the B-cell response within the liver.

### The transcriptomic signature of liver-autoreactive CD4 T cells is dictated by the hepatic environment

Since circulating liver-autoreactive CD4 T cells have a specific transcriptomic signature in humans, we investigated whether their molecular profile was due to antigen reactivity in the hepatic environment or to a lineage-specific signature. To answer this question, we designed a non-TCR-transgenic mouse model that allows the analysis of the emergence and modulation of CD4 T cell responses against a model antigen expressed by hepatocytes. We used a model of Balb/c mice in which the hemagglutinin (HA) expression was restricted to hepatocytes, by an inducible Cre recombinase under the control of hepatocyte-specific mTTR (transthyretin) promotor (HA/iCre) (Figure 6A and Supplementary Figure 11). Tamoxifen induced HA expression in the liver of HA/iCre mice but did not induce an anti-HA humoral response (Figures 6A-C). Intramuscular injection of a Cre-coding adenovirus (AdCre) or an HA-coding plasmid was performed to generate peripheral immunization against the HA antigen (HA i.m, Figure 6A and C). HA-specific memory (CD44^high^) CD4 T cells were detected with pMHCII tetramer in the spleen but not in the liver of mice after peripheral immunization.

**Figure 6:**
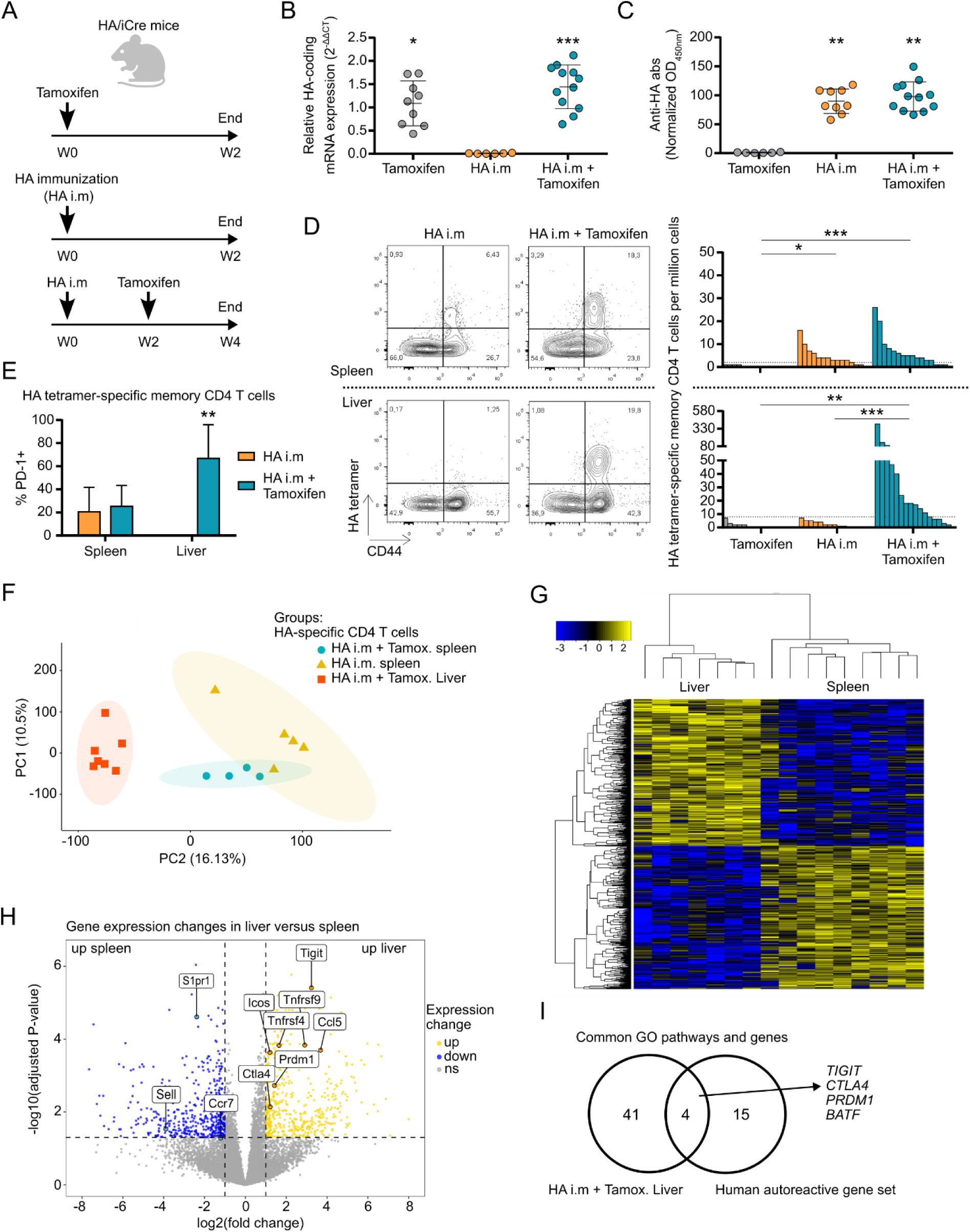
Analysis of antigen-specific CD4 T cells from the liver and the spleen of an in vivo non-TCR-transgenic mouse model. (A) Experimental design for investigation of immune response against the HA antigen in the liver. HA/iCre mice received either tamoxifen treatment (n=9), HA immunization treatment (HA i.m; n=15), or HA immunization followed 2 weeks later by tamoxifen treatment (HA i.m + Tamoxifen; n=12). Mice were euthanized 2 weeks after the last treatment. (B) Relative HA mRNA expression in the liver. ACTB was used as loading control. (C) Analysis of normalized anti-HA antibody rate in serum of mice. (D) Analysis of HA-tetramer-specific CD4 T cells in the spleen (top) and in the liver (bottom). Contour plot representation of HA tetramer staining and CD44 expression in CD4 T cells from HA i.m and HA i.m + Tamoxifen mice (left). Frequency of HA-tetramer-specific memory (CD44^high^) CD4 T cells per million cells (right). Dotted lines in the frequency graphs represent threshold of positive detection of HA-specific CD4 T cell population. (E) Analysis of PD-1 expression in detectable HA-specific memory CD4 T cells from the spleen and the liver of HA i.m and HA i.m + Tamoxifen mice. (F) Principal component analysis (PCA) of bulk-RNA-seq transcriptome of HA-specific CD4 T cells isolated from the liver or the spleen. (G) Gene expression heatmap of HA-specific CD4 T cells. (H) Volcano plot of gene expression change between liver and spleen HA-specific CD4 T cells. (I) Genes from GO enrichment pathways upregulated and shared between mouse liver HA-specific CD4 T cells and human autoreactive CD4 T cells. Dunn’s multiple comparisons test was used for B, C and D. Mann-Whitney test was used for E. *: p<0.05; **: p<0.01; ***: p<0.001.

To analyze HA-specific CD4 T cells in the liver, we induced HA expression in the liver, by using tamoxifen in pre-immunized HA/iCre mice (HA i.m + Tamoxifen, Figure 6A); we used littermates lacking liver-restricted expression of an inducible Cre recombinase (Ctrl) as controls (Supplementary Figure 11). In this condition, we detected HA-specific CD4 T cells in the liver of immunized mice expressing HA by hepatocytes after tamoxifen treatment (Figure 6D), showing active recruitment from the periphery of HA-specific memory CD4 T cells into the liver following local antigen expression. Liver HA-specific CD4 T cells expressed high levels of PD-1 (Figure 6E).

In this mouse model, we analyzed the transcriptomic profile of HA-specific CD4 T cells in mouse liver and spleen. Overall, all spleen HA-specific CD4 T cells, irrespective of liver HA expression, clustered together, whereas liver HA-specific CD4 T cells show a distinct transcriptomic signature (Figure 6F and G). Besides expressing genes associated with the inflammatory response (e.g. *Ox40*, *Cd137*, *Ccl5* and *Icos*), liver HA-specific CD4 T cells expressed genes involved in the regulation of proliferation and activation (e.g. *Tigit*, *Ctla4*, *Prdm1* and *Casp3*) (Figures 6H). Comparative analysis with human liver-self-antigen specific transcriptomic signature revealed four common pathways linked to the regulation of T cell activation (GO:0042110 T cell activation; GO:0002694 reg. of leukocyte activation; GO:0051249 reg. of lymphocyte activation; GO:0050863 reg. of T cell activation). The analysis of the genes from these pathways revealed common genes shared between liver HA-specific CD4 T cells in mice and circulating liver-autoreactive CD4 T cells in human (Figure 6I). Both cell types expressed high levels of *TIGIT*, *CTLA-4, BATF* and *PRDM1* gene expression. These data demonstrate immuno-regulatory genes were induced during local immune response in the liver. These suggest that the immune signature of human circulating liver-autoreactive CD4 T cells is an imprint of the hepatic environment, and could suggest that these cells leak out of damaged tissue after a phase of high reactivity in the tissue.

### Immune checkpoint molecules control the antigen-specific hepatitis in a CD4 T cell dependent manner

The upregulation of immune checkpoint molecules by the antigen-specific CD4 T cells after their reactivity in the liver suggested the potential regulation of these cells by the hepatic environment. Therefore, we blocked PD-1 and CTLA-4 pathways, with specific antibodies as immune checkpoint inhibitors (ICI), in our mouse model of hepatic infiltration by HA-specific CD4 T cells, after immunization and induction of HA expression by hepatocytes (Figure 7A). The obtained data revealed a significant hepatitis marked by high histological liver inflammation score when mice receiving ICI were pre-immunized and expressed HA in the liver (HA/iCre + ICI, Figures 7B and C). Mice immunized but unable to express HA in their liver after tamoxifen treatment did not show significant liver inflammation after PD-1 and CTLA-4 blockade (Ctrl + ICI, Figures 7B and C). These data demonstrate that the hepatic environment specifically regulates the antigen-specific response by engaging immune checkpoint receptors.

**Figure 7:**
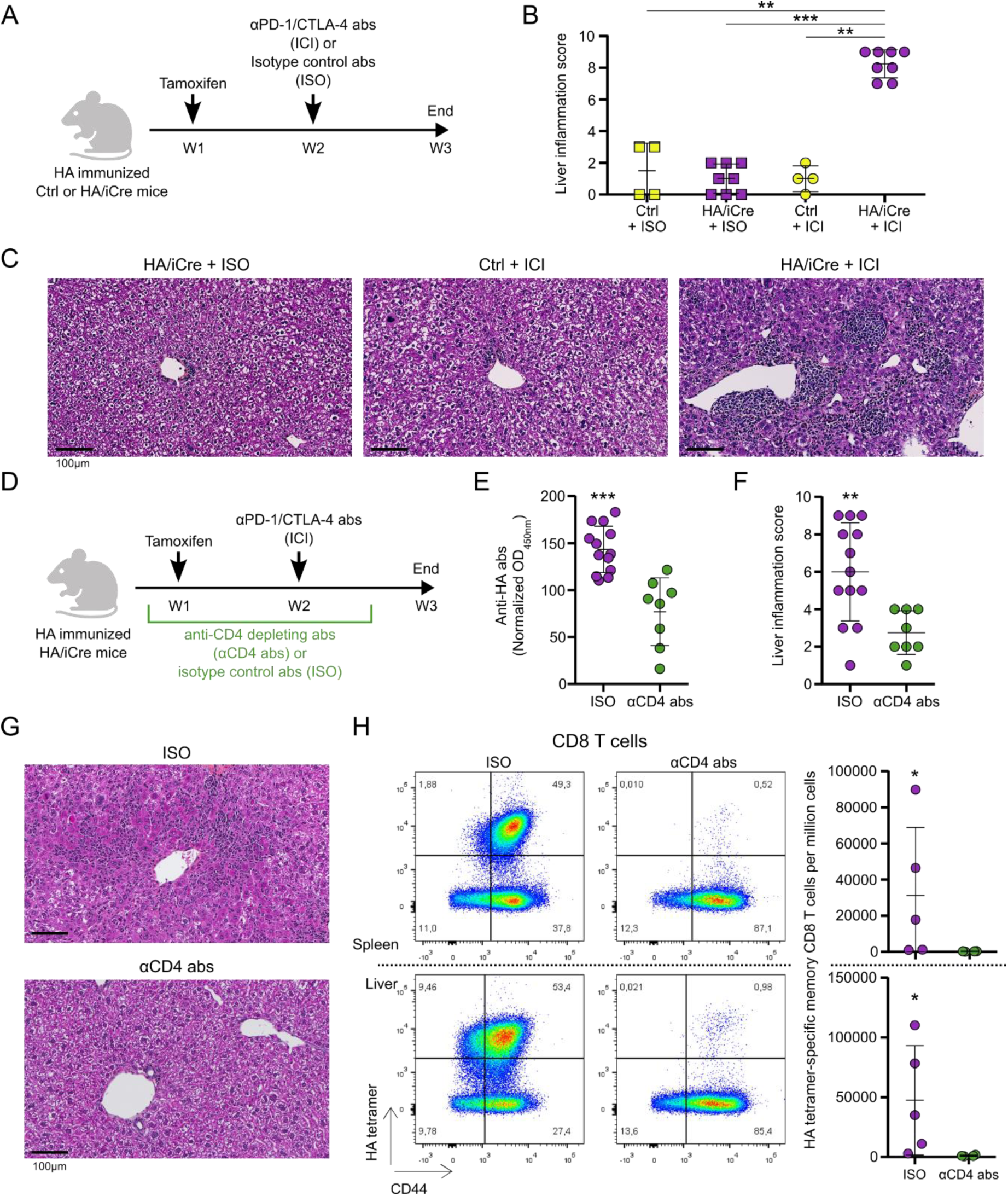
Immune checkpoint pathways control liver antigen-specific CD4 T cell responses and mediate hepatic tolerance. (A) Experimental design for PD-1 and CTLA-4 blockade (ICI) in HA immunized tamoxifen-treated HA/iCre mice. Injection of isotype control antibodies to ICI was used as control (ISO). (B) Histological liver inflammation scoring analysis of liver tissue sections from immunized tamoxifen-treated control (Ctrl; n=8) and HA/iCre (n=16) mice treated either with isotype control antibodies (+ ISO; Ctrl: n=4, HA/iCre: n=8) or anti-PD-1 and anti-CTLA-4 blocking antibodies (+ ICI; Ctrl: n=4, HA/iCre: n=8). (C) Representative pictures of paraffin-embedded liver sections stained with HPS coloration from indicated conditions. Black line is used as scale. (D) Experimental design for CD4 depletion (αCD4 abs) in immunized tamoxifen-treated HA/iCre mice receiving PD-1 and CTLA-4 blockade protocol (n=13). Injection of isotype control antibody to αCD4 abs was used as control (ISO; n=8). (E) Analysis of normalized anti-HA antibody rate in serum of mice. (F) Histological liver inflammation scoring analysis of liver tissue sections. (G) Representative pictures of paraffin-embedded liver sections stained with HPS coloration from indicated conditions. Black line is used as scale. (H) Pseudocolor dot plot representation of HA tetramer staining and CD44 expression in CD8 T cells from spleen (top) and liver (bottom) of ISO (n=5) and αCD4 abs (n=4) mice (left). Frequency of HA tetramer-specific memory (CD44^high^) CD8 T cells per million cells (right). Mann-Whitney test was used for B, E, F and H. *: p<0.05; **: p<0.01; ***: p<0.001.

Then, we assessed the role of CD4 T cells in this response by depleting CD4^+^ cells with anti-CD4 antibodies in immunized HA/iCre mice treated with tamoxifen and PD-1/CTLA-4 blocking antibodies (Figure 7D). First, CD4 depletion was associated with significantly lower immunization rate according to anti-HA antibody levels in the serum (Figure 7E). Interestingly, histological analysis revealed significantly reduced liver damages in CD4 depleted mice, showing major role of CD4 T cells in promoting antigen-specific liver inflammation in our model (Figures 7F and G). The hepatic inflammation observed after PD-1 and CTLA-4 blockade was linked to substantial HA-specific CD8 response in the spleen and in the liver, but depletion of CD4 T cells before the induction of HA expression in the liver abrogated the HA-specific CD8 response (Figure 7H). These results demonstrate that the hepatic environment quickly regulates local immune response against an antigen expressed by hepatocytes through upregulation of immune checkpoint pathways in a CD4-dependent manner. This data reveal a major role of the liver-antigen-specific CD4 T cells in the initiation of immune response against a hepatic antigen.

## Discussion

In this study, we analyzed the transcriptome, TCR repertoire and single-cell protein expression of circulating liver-self-antigen-specific CD4 T cells and expanded intrahepatic CD4 T cell clonotypes found in the blood of AILD patients. We have identified a peripheral subset of liver-autoreactive CD4 T cells (non-T_REG_ T_CM_ CXCR5^-^ CD49d^+^ PD-1^+^ TIGIT^+^ ICOS^+^ HLA-DR^+^ CD38^+^ CD4 T cells) with an autoimmune transcriptomic profile (*CXCL13, IL21, TIGIT, CTLA4, PRDM1*, *TOX, HLA-DR, CCR2, ENTPD1, LGALS1, CXCR3, CXCR6, TIM3*) that is linked to clonal expansion in the liver. Using a mouse model to track the early immune response against an antigen conditionally overexpressed in the liver, we observed that liver CD4 T cells targeting this antigen acquired an immuno-exhausted (*TIGIT, PRDM1, CTLA-4)* transcriptomic signature comparable to that of circulating liver-autoreactive CD4 T cells in AILD patients, and are essential for inducing a complete immune response against a liver antigen responsible for liver damage. Our data support the idea that circulating CD4 T cells targeting hepatic antigens are liver-derived autoreactive CD4 T cells and that their transcriptomic signature is imprinted by antigen encounter in the tissue. The ability to identify a subset of T cells linked to tissue auto-reactivity in blood opens important prospects for the development of biomarkers and therapies. This is in line with the capacity to identify tumor-specific CD8 T cells in blood which shares similarities with their counterparts in tissues, and could therefore be an alternative source for identifying antitumor T cell reactivity^37^.

This study also represents the first complete deep phenotyping and transcriptome analysis of CD4 T cells targeting hepatic autoantigens in human AILD. We analyzed the transcriptomic signature of the CD4 T cell reactivity against three distinct liver-self-antigens and against three common foreign antigens. Upon *in vitro* peptide stimulation, autoreactive CD4 T cells present a common transcriptomic signature although the three self-antigens have no molecular similarity (Sepsecs, CYP2D6 and PDCE2). Interestingly, this transcriptomic signature is distinct from the anti-fungus Th17 (Candida Albicans reactivity) and the anti-viral (influenza H1N1 and Sars-CoV-2) immune signatures. In a previous publication, we had shown that Sepsecs-specific CD4 T cells, versus CD4 T cells reactive against the Candida Albicans, have a B-helper / pro-inflammatory immune signature^22^. In this new study, we reinforce the data, and we demonstrate that the immune signature of liver-autoimmunity is clearly distinct from diverse foreign antigen immune responses. The major immune genes characterizing this signature are *TIGIT, CTLA4, NR3C1, TOX, CXCL13, STAT3,* and *PRDM1*. This autoreactive signature shares a high similarity with the published signature of the T_PH_ CD4 cell subset described in the tissue of other autoimmune diseases, ^23,24,29,38^. However, the lack of data available at the single-cell level for other self-antigen-specific CD4 T cells after short peptide stimulation makes it difficult to identify genes specifically linked to hepatic autoimmunity in our dataset.

Interestingly, circulating liver-autoreactive CD4 T cells have a profile similar to that of tissue-infiltrating CD4 T cells found in various immune contexts. Expression of PD-1, CXCL13, IL21, TIGIT, PRDM1, TOX, CD39, TIM3, CTLA-4, CXCR3, CCR2, HLA-DR and CXCR6 is commonly shared with the signature of T_PH_ CD4 cells in the synovium of RA patients, neoantigen-reactive CD4 T cells in metastatic human cancers, tumor-infiltrating CD4 T cells, liver-infiltrating CD4 T cells in HBV-infected patients and liver-resident CD4 T cells^23,39–46^. The main difference is that circulating liver-autoreactive CD4 T cells do not express CD69, which is probably downregulated after tissue egress. This supports the idea that local antigenic reactivity induces a significant modification of CD4 T cells, characterized by the induction of the expression of immune checkpoint molecules and other. These markers found in the peripheral blood may be useful for designing biomarkers related to tissue aggression. Their presence in the blood stream can be explained by the fact that during the active phase of the disease, the high clonal expansion in the liver may favor the recirculation of autoreactive T cells. This concept is supported by the fact that the frequency of liver-autoreactive CD4 T cells in the blood of patients decreases during the remission phase, which is generally associated with a reduction in intrahepatic lymphocyte activity.

The most common feature shared between liver-autoreactive CD4 T cells and tissue infiltrating CD4 T cells is their high expression of immune-regulatory molecules (e.g. PD-1, TIGIT, CTLA-4), which may be linked to local chronic antigen stimulation and exhaustion. During cancer, the blockade of PD-1 can restore the activity of tumor-infiltrating CD4 T cells demonstrating their local immune involvement and that immune checkpoint molecules control local reactivity^41,43^. In our study, we demonstrate that immune checkpoint molecules in the liver participate in the local control of antigen-specific liver damages which is dependent of the CD4 T cell response. In human, hepatic lesions can be induced by anti-cancer immunotherapy (anti-PD-1 and anti-CTLA-4) and are associated with a predominant CD8 T cell infiltrate in the lobular zone, suggesting a reactivation of the local liver-antigen-specific T cell response^47–49^. The major difference with AIH is that during immunotherapy-induced acute hepatitis, no signs of chronicity are reported, and the acute hepatitis is rapidly controlled by immunosuppressive treatments without relapse after treatment withdrawal. In contrast, AIH patients’ liver biopsies show the complete architecture of a chronic adaptive immune response at diagnosis, which is usually late in the course of the disease. A large number of B cells and of CD4 T cells are observed in the portal zone^48,49^. The detection of tertiary lymphoid structures in the portal zone in our study supports the idea of a prolonged activation period to achieve such immune organization.

These differences between acute and chronic hepatitis suggest a possible gradual escape from standard immuno-regulatory processes of the liver-self-antigen-specific T cells during AILD. Studies have shown links between CTLA-4 polymorphisms and susceptibility to AIH and PBC^50–55^. However, the functional biological link with the development of AILD is not known. We can hypothesize that a dysregulation of this pathway participates in the gradual temporal escape of liver-antigen-specific CD4 T cell response that mediates the expansion of cytotoxic CD8 T cells. This hypothesis may be supported by the fact that we were able to detect persistent self-antigen-specific CD4 T cell clones for three years, suggesting maintenance of clonal expansion. Therefore, understanding the link between genetic risk and the functional capacity of autoreactive T cells to escape tolerance may open new therapeutic prospects.

## Methods

### Patients

All the patients eligible signed a written informed consent prior to inclusion into a bio-bank of samples of AILD patients (BIO-MAI-FOIE) maintained in Nantes University Hospital which obtained regulatory clearance from the biomedical research Ethics Committee (CPP) and all required French Research Ministries authorizations to collect biological samples (ref MESR DC-2017-2987). The biobank is supported by the HEPATIMGO network promoted since 2017 (RC17_0228) by Nantes University Hospital and is a prospective multi-centric collection managed by the Biological Resource center of the CHU of Nantes. All data collected were treated confidentially and participants are free to withdraw from the study at any time, without explanation and without prejudice to their future care. It was granted authorization from the CNIL: 2001209v0). All AIH patients included in this study had a simplified diagnostic score superior or equal to 6 according to the simplified scoring system for AIH of the international autoimmune hepatitis group (IAHG)^5,56^. PBC patients included in this study were diagnose based on cholestasis, increase serum IgM and presence of anti-mitochondria M2 (anti-M2) antibody^7^. Active AIH (AIHa) patients are untreated patients with new onset AIH patients enrolled at diagnosis prior any treatment initiation as previously described^57^, and AIH patients under standard treatment but do not normalize the transaminases (AST and ALT) and/or the serum IgG levels or are under relapsing event. Remission AIH (AIHr) patients are defined biochemically by a normalization of the transaminases and the IgG levels, according to the most recent European clinical practice guidelines on AIH ^5^. All NASH (non-Alcoholic SteatoHepatitis) patients had histological evidence of NASH and had dysmetabolic syndromes. This study was carried out in accordance with the Principles of International Conference on Harmonisation (ICH) Good Clinical Practice (GCP) (as adopted in France) which builds upon the ethical codes contained in the current version of the Declaration of Helsinki, the rules and recommendations of good international (ICH) and French clinical practice (good clinical practice guidelines for biomedical research on medicinal products for human use) and the European regulations and/or national legislation and regulations on clinical trials.

### Peptide re-stimulation assay

10 to 20×10^6^ PBMCs (at a final concentration of 10×10^6^/mL) were stimulated for 3 to 4h at 37°C with 1µg/mL of synthesized peptides spanning all of the Sepsecs, CYP2D6 or PDCE2 sequences (20 amino acids in length with a 12 amino acid overlap; Synpeptide, China) or with 1µg/mL of PepTivator^R^ C. albicans MP65, Influenza A (H1N1) MP1 and SARS-CoV-2 Prot_S (SPIKE) (peptides pools of 15 amino acids length with 11 amino acid overlap, Miltenyi Biotec) in 5% human serum RPMI medium in the presence of 1µg/ml anti-CD40 (HB14, Miltenyi Biotec). After 4 hrs of specific peptide stimulation, PBMCs were first labeled with PE-conjugated anti-CD154 (5C8, Miltenyi Biotec) and CD154^+^ cells were then enriched using anti-PE magnetic beads (Miltenyi Biotec). A 1/10th fraction of non-enriched cells was saved for frequency determination. Frequency was calculated with the formula F = n/N, where n is the number of CD154 positive cells in the bound fraction after enrichment and N is the total number of CD4^+^ T cells (calculated as 10 × the number of CD4^+^ T cells in 1/10th non-enriched fraction that was saved for analysis). After enrichment, cells were stained with appropriate antibodies (Supplementary table 1).

### Tetramer staining

For tetramer staining, 30 million PBMCs in 200 µL of 5% human serum RPMI medium were stained with 20 µg/mL PE-labeled tetramers at room temperature for 100 minutes. Cells were washed and incubated with anti-PE magnetic beads (Miltenyi Biotec, Germany), and a one-tenth fraction was saved for analysis. The other fraction was passed through a magnetic column (Miltenyi Biotec). After enrichment, cells were stained with appropriate antibodies (Supplementary table 1). Sepsecs_187-197_ HLA-DRB1 03:01 tetramer was generated by the tetramer core at the Benaroya research institute (Seattle, USA).

### Flow cytometry and cell sorting

All antibodies used are described in the supplementary table 1. Briefly, for surface staining, PBMCs were incubated 20 minutes with a mix of antibodies and then washed prior analysis or cell sorting on BD FACSCantoII, Cytek Aurora or BD FACSAriaII. For intra-cellular staining we used the Fixation/Permeabilization Solution Kit (BD Cytofix/Cytoperm™, BD Biosciences).

### Single cell RNA sequencing and analysis (plate-based assays)

For scRNA-seq analysis of peptide-restimulated CD4 T cells, CD154^+^ memory CD4 T cells were first sorted on BD FACSAriaII, one cell per well, in 96-well plates containing specific lysis buffer at the CRTI, Nantes. Plates were immediately frozen for storage at −80°C, and sent on dry ice to the Genomics core facility of CIML, Marseille, for further generating scRNAseq libraries with the FB5P-seq protocol as described^22,58^. Briefly, mRNA reverse transcription (RT), cDNA 5’-end barcoding and PCR amplification were performed with a template switching (TS) approach. After amplification, barcoded full-length cDNA from each well were pooled for purification and library preparation. For each plate, an Illumina sequencing library targeting the 5’-end of barcoded cDNA was prepared by a modified transposase-based method incorporating a plate-associated i7 barcode. Resulting libraries had a broad size distribution, resulting in gene template reads covering the 5’-end of transcripts from the 3^rd^ to the 60^th^ percentile of gene body length on average. As a consequence, sequencing reads covered the whole variable and a significant portion of the constant region of the TCRα and TCRβ expressed mRNAs, enabling assembly and reconstitution of TCR repertoire from scRNAseq data. Libraries prepared with the FB5P-seq protocol were sequenced on Illumina NextSeq2000 platform with P2 100-cycle kits, targeting 5×10^5^-1×10^6^ reads per cell in paired-end single-index mode with the following configuration: Read1 (gene template) 103 cycles, Read i7 (plate barcode) 8 cycles, Read2 (cell barcode and Unique Molecular Identifier) 16 cycles. We then used a custom bioinformatics pipeline to process fastq files and generate single-cell gene expression matrices and TCR sequence files, as described^22,58^.

For scRNA-seq of tetramer-stained CD4 T cells, we used an updated version of FB5P-seq (Flash-FB5P-seq) modified to implement advances developed by Hahaut et al. in the Flash-seq protocol^59^. Cells were sorted in 2 µl lysis mix composed of 0.1% Triton X-100 (0.02 µl), 1.2 U/µl recombinant RNAse inhibitor (0.06 µl), 6 mM dNTP mix (0.48 µl), 1.6 mM RT primer (5’-TGCGGTATCTAAAGCGGTGAGTTTTTTTTTTTTTTTTTTTTTTTTTTTTTT*V*N-3’) (0.36 µl), 0.0125 pg/µl ERCC spike-in mix (0.05 µl), 9 mM dCTP (0.18 µl), 1 M betaine (0.4 µl), 1.2 mM DTT (0.024 µl) and PCR-grade H_2_O (0.426 µl). Plates were conserved at −80°C until sorting, thawed at RT for cell sorting, and immediately refrozen on dry ice then conserved at −80°C until further processing. Reverse transcription with template switching and cDNA amplification were performed in a single step. Each plate containing cells in lysis mix was thawed on ice, heated at 72°C for 3 min then immediately placed on ice. To each well containing 2 µl cell lysate, we added 8 µl of an RT-PCR mix composed of 2 U/µl Superscript IV reverse transcriptase (0.1 µl), 0.8 U/µl recombinant RNAse inhibitor (0.2 µl), 1X KAPA HiFi ReadyMix (5 µl), 4.8 mM DTT (0.48 µl), 0.8 M betaine (1.6 µl), 9.2 mM MgCl_2_ (0.092 µl), 0.1 µM forward PCR primer (5’-AGACGTGTGCTCTTCCGAT*C*T-3’) (0.1 µl), 0.1 µM reverse PCR primer (5’-TGCGGTATCTAAAGCGGTG*A*G-3’) (0.1 µl), 1.84 µM biotinylated barcoding template switching oligonucleotide (5’-biotin-AGACGTGTGCTCTTCCGATCTXXXXXXXXNNNNNNNNCAGCArGrGrG-3’, where XXXXXXXX are well-specific barcodes as published^58^, NNNNNNNN are random nucleotides serving as Unique Molecular Identifiers, and rG are riboguanosine RNA bases) (0.184 µl) and PCR-grade H_2_O (0.144 µl). Then plates were incubated in a thermal cycler 60 min at 50°C, 3 min at 98°C, and 22 cycles of 20 sec at 98°C, 20 sec at 67°C and 6 min at 72°C. We then pooled 5 µl amplified cDNA from all wells into one tube per plate, and performed 0.6X SPRI bead-based purification. Sequencing libraries were prepared from 800 pg cDNA per plate with the Illumina Nextera XT protocol, modified to target sequencing reads to the 5’ end of cDNA, as described ^22,58^. Before sequencing, pooled indexed libraries were depleted from ribosomal RNA-derived cDNA molecules by using the SEQoia RiboDepletion Kit (Biorad) according to the manufacturer’s instructions. Flash-FB5P-seq libraries were sequenced on an Illumina NextSeq2000 using P3 100 cycle kit and targeting 5×10^5^-1×10^6^ reads per cell in paired-end single-index mode with the following configuration: Read1 (gene template) 114 cycles, Read i7 (plate barcode) 8 cycles, Read2 (cell barcode and Unique Molecular Identifier) 16 cycles. We then used a custom bioinformatics pipeline to process fastq files and generate single-cell gene expression matrices and TCR sequence files. Briefly, reads were processed with the zUMIs pipeline^60^ to generate the gene expression UMI count matrix, and with the TRUST4 pipeline^61^ to reconstruct TCR sequences.

Quality control excluded cells with more than 10% mitochondrial UMIs, less than 400 genes, more than 25% ERCC UMIs, less than 0.7 ERCC spike-in quantification accuracy^58^, and less than 5% ribosomal protein coding UMIs. Gene expression UMI matrices were analyzed with the *Seurat v4.1.0* package in R v4.1.2 using standard log normalization, 4,000 highly variable genes (computed with the *vst* method) from which we removed all TCR coding genes, 40 principal components for computing UMAP and Louvain clustering at resolution 0.8. Marker genes of unsupervised clusters or metadata-defined clusters were computed with the *FindAllMarkers* function using Wilcoxon test with an adjusted p-value cutoff of 0.01. Gene expression heatmaps were generated with the *pheatmap* function. TCR clonotypes were computed and analyzed as described previously^22^.

### Single cell RNA sequencing and analysis (10X Genomics protocol)

First, PD-1^+^CD45RA^-^CXCR5^-^ and PD-1^-^CD45RA^-^CXCR5^-^ CD4 T cells were sorted on BD FACSAriaII. Cells were counted, centrifuged (500g, 5 min, 4°c) then resuspended at the recommended dilution for a 20000 cell loading per sample onto a Next GEM Chip K (PN-1000286) and run on a Chromium Single Cell Controller using the Chromium Single Cell 5′ V2 Next GEM single cell kit (PN-1000263) according to the manufacturer’s instructions (CG000331, 10X Genomics). Gex and VDJ-TCR libraries were then prepared (PN-1000190, PN-1000252, PN-1000215 also from 10X genomics), checked for quality controls and sequenced on a S2 flow cell on a Nova-Seq 6000 (Illumina) at the GenoBird platform (IRS-UN, CHU Nantes). Raw reads were analyzed using FastQC for quality controls and were then processed using CellRanger pipeline (v3.1.0 with default parameters). Generated FASTQ files were aligned to the human reference genome GRCh38.

The base calling has been done using Illumina bcl2fastq2 (v2.20). The FASTQs obtained has been analyzed for cell identification and UMIs count using 10X Genomics Cell Ranger 7.0.1 and hg38 reference. Four filtered matrices were obtained for each sample and the script written for their analysis has been written in R (v4.2.2). The matrices have been analyzed using Seurat 4^62^ using default parameters. Each matrix has been filtered with the following cut-off: 500 < Features_RNA < 4000; percent.mt < 5 and cells having two distinct TRA and TRB identified have been removed for the rest of the analysis. Then each matrix has been normalized using the SCTransform method with a regression on the percentage of ribosomal genes count. The four matrices has been merged and integrated using harmony package^63^. The cell clusters have been identified using harmony for reduction. The markers for each clusters have been identified with a min.pct=0.25 (genes detected in a minimum fraction of min.pct cells) and a threshold of logFoldChange of 0.25. The markers retained for each cluster where those having a p-value adjusted < 0.01. The annotation of clusters has been done using azimuth package^62^. The module score has been calculated for each cluster using a predefined list of genes for “Auto-reactive”, “SARS-CoV-2”, “H1N1” and “C.ALB”. The top five percent of cells base on this module score has been highlighted on the UMAP. The clusters have been then compared based on their module score for each gene list using Kruskal Wallis test followed by a paired Wilcoxon rank test.

### Mini-bulk-RNA sequencing and analysis

For mini-bulk RNA-seq of *in vitro* stimulated CD4 T cells, we used a slightly modified version of the Flash-FB5P-seq protocol described above. First, CD4 T cell subsets were sorted on BD FACSAriaII. 500 to 2000 cells per well were stimulated in presence of 2.5µg/ml anti-CD3 and 1µg/ml anti-CD28. After 24h, cells were washed gently in PBS and resuspended in 4µL of specific lysis buffer with the composition described above except for ERCC spike-in mix which we used 10-time more concentrated (0.125 pg/µl). We performed RT-PCR with 16 µl of the RT-PCR mix described above, using 16 cycles of PCR for cDNA amplification (instead of 22 cycles for single-cell assay). Sequencing libraries were prepared and sequenced as described above for Flash-FB5P-seq. Sequencing reads were processed with the zUMIs pipeline^60^ to generate gene expression UMI count matrices. Mini-bulk gene expression UMI count matrices were pooled across plates and further processed with the *DEseq2* package for normalization (*vst* method) and computation of differentially expressed genes between non-stimulated and TCR-stimulated cells from each sorted phenotype. The mean of normalized expression values for manually selected significant DE genes was used to construct the heatmaps in Figure 5F.

### Bulk TCRβ sequencing

100×10^3^ PD-1^+^CD45RA^-^CXCR5^-^ and PD-1^-^CD45RA^-^CXCR5^-^ CD4 T cell subsets were sorted on a BD FACSAriaII. gDNA from T cells and frozen liver biopsies was extracted with NucleoSpin® Blood kit (Macherey-Nagel) and TCRβ sequencing was performed as previously described^64^.

### Unsupervised flow cytometry analysis

Flow cytometry data from patients were analyzed using Omiq software (Dotmatics, USA). Subsampling of viable lymphoid CD45RA^-^ CD4^+^ CD3^+^ cells was performed (20×10^3^ per patients). Clustering was performed using FlowSOM to generate 150 clusters. 22 Metacluster were generated by running consensus metaclustering. Wilcoxon test was used to identify significant variation between two groups of patients.

### Spatial multi-phenotyping and analysis (Phenocycler instrument)

The PhenoCycler^TM^ instrument (Akoya Biosciences, USA) performs iterative annealing and removal of fluorophore-conjugated oligo probes to primary antibody-conjugated complementary DNA barcodes. Antibody panel was constructed using 11 ready-to-use commercially available PhenoCycler^TM^ antibodies (Akoya Biosciences) and one antibody unavailable has been custom conjugated. Detailed 12-plex PhenoCycler^TM^ information is provided in Supplementary table 1. Anti-CXCR5 antibody has been custom conjugated with a unique PhenoCycler^TM^ oligonucleotide tag using the antibody conjugation kit of Akoya Biosciences following their instructions.

FFPE human liver was sectioned to a thickness of 5 µm and directly adhered onto poly-L-lysine (Sigma Aldrich) coated 22 x 22 mm coverslips (Akoya Biosciences). Tissue coverslips were stored at 4°C until staining. The sample was incubated for 20 minutes at 55°C on a hot plate and tissue was cooled down before staining. Tissue section was deparaffinized and rehydrated by immersing the coverslip through the following solution for 5 minutes each: twice in OTTIX, twice in 100% Ethanol, once in 90%, 70%, 50%, 30% Ethanol and twice in ddH2O. Antigen retrieval was performed in a hot water bath at 100°C in a 1X Citrate Buffer (Sigma Aldrich) for 30 minutes. After cooling at room temperature, tissue section was briefly washed twice in ddH2O for 2 minutes. Following hydration step, tissue section was blocked in Staining buffer at room temperature for 20 minutes and stained with the 12 plex PhenoCycler^TM^ antibody panel in a humidity chamber 3 hours at room temperature. To ensure that the antibodies remain attached to the antigen during the multicycles PhenoCycler^TM^ imaging, protocol post-staining protocol has 3 fixing steps. After incubation with antibodies, tissue section was washed twice and post-fixed with 1.6% PFA Storage buffer for 10 minutes at room temperature. Tissue sections were briefly washed three times in 1X PBS buffer and incubated in a cold methanol solution for 5 minutes. Following washing, tissue section was fixed in final fixative solution for 20 minutes at room temperature. Finally, tissue section was washed three times in 1X PBS buffer and was stored in Storage Buffer at 4°C before PhenoCycler^TM^ imaging.

PhenoCycler^TM^ imaging was performed with an Axio Observer (Zeiss) inverted microscope equipped with camera ORCA Flash 4.0 LT (Hamamatsu). The microscope is coupled to the Colibri 7 LED light source (Zeiss) and fluorescence was detected using the monoband filter Set 112 HE LED (Zeiss). The PhenoCycler^TM^ experimental run was managed by the instrument controller software (v1.30.0.12, Akoya Biosciences) integrating with the Zeiss microscope. Nuclear DAPI staining was used to design manually tiled regions of interest at the 5x magnification (N-Achro 5x/0.15 M27, Zeiss). Automated multiplex imaging was performed using a Plan Apo 20x/0.8 M27 Air objective (Zeiss) with a 325 x 325 nm pixel size using software autofocus repeated every tile before acquiring a 11 plane-z-stack with a z-spacing of 1.5 µm. Data are acquired as 3D stacks by cycle of 4 markers (including a recurrent nuclear DAPI staining). After acquisition, raw files were exported using the CODEX Instrument Manager (CIM, Akoya Biosciences). The data are processed by computing the more in focus image from the stack to take into account the potential tissue unflatness, every cycles are registered together based on the DAPI staining, and the fluorescence signal is corrected by background fluorescence based on blank cycles and normalized in intensity. Cell segmentation was done by using Cellpose algorithm. After segmentation, measurement data are transformed in FCS files and analyzed with the CODEX MAV software.

### Mice and treatments

Heterozygous transthyretin (TTR)-inducible Cre (iCre) mice, originally on a C57Bl6 backgroud^65^, were back-crossed on a Balb/c background for at least 10 generations (TAAM, CDTA CNRS Orléans, FRANCE). They were cross-bred with homozygous Rosa26 hemagglutinin (HA) floxed mice mice (Rosa26^tm(HA)1Libl^, Kindly provided by R. Liblau, Toulouse, France^66^), resulting in heterozygous Rosa26 HA floxed (Ctrl) mice and Rosa26 HA floxed TTR-inducible Cre (HA/iCre) mice. Mice were genotyped for Cre-coding gene at 3 weeks-old thanks to small tail biopsies using following Cre primers: 5’-CCTGGAAAATGCTTCTGTCCG-3’ sense sequence and 5’-CAGGGTGTTATAAGCAATCCC-3’ antisense sequence. Male and female eight to twelve-weeks-old mice were used for each experiment. All mice were housed at the UTE IRS-UN animal facilities (Nantes, FRANCE) where they were fed *ad libitum* and allowed continuous access to tap water. Procedures were approved by the regional ethical committee for animal care and use and by the Ministère de l’enseignement supérieur et de la recherche (agreements APAFIS #2054, #28582 and #43529). All experiments were performed in accordance with relevant guidelines and regulations.

Induction of HA expression by hepatocytes was performed by feeding mice with tamoxifen dry food (0.5g/kg tamoxifen + 5% saccharose; Safe, FRANCE) for 14 days in free access or by injecting intraperitonially (i.p) 1mg of tamoxifen diluted in corn oil three times in two weeks span. Peripheral immunization against HA was performed with two different materials: one intramuscular (i.m) injection of 1,5.10^9^ infectious particle of adenoviral CAG Cre vectors (AdCre, produced by INSERM UMR 1089 CPV facility, Nantes, FRANCE) in *tibialis anterior* of one posterior leg or i.m injection of 50µg of plasmid CMV HA vector in *tibialis anterior* of both posterior legs, twice three weeks apart.

Immune checkpoint blockade protocol was performed by intravenous (i.v) injection of 200µg of anti-mouse PD-1 antibody (29F.1A12™ clone, BioXCell) and 200µg of anti-mouse CTLA-4 antibody (9H10 clone, BioXCell), using purified Rat IgG2a,κ isotype control antibody (RTK2758 clone, Biolegend) and polyclonal Syrian Hamster IgG antibody (ref BE0087, BioXCell) as controls, three times in one week span. Depletion of CD4^+^ cells was performed by i.p injection of 500µg of anti-mouse CD4 antibody (GK1.5 clone, BioXCell), using purified Rat IgG2b,κ isotype control antibody (LTF-2 clone, BioXCell), every two days during two weeks.

### RNA extraction, reverse transcription and quantitative PCR

Total RNA was extracted from organ tissue using TRIzol^TM^ reagent (15596026, ThermoFisher Scientific) and purified with RNeasy Mini Kit (74106, Qiagen) according to the manufacturer protocol. Reverse transcription was performed using 2µg of total RNA incubated at 70°C for 10min with poly-dT24 20µg/mL (Eurofins Genomics), 8mM DTT (18057018, ThermoFisher Scientific) and 20mM of each dNTP (10297018, ThermoFisher Scientific). After a brief 5min incubation at 4°C, first strand buffer 5X (Y00146, Invitrogen), 200U of M-MLV reverse transcriptase (18057018, ThermoFisher Scientific) and 40U of RNAse OUT inhibitor (10777019, Invitrogen) were added and incubated at 37°C for 1h followed by 15min at 70°C. Real-time quantitative PCR was performed using the ViiA^TM^ 7 Real-Time PCR System and Power SYBR^TM^ Green PCR Master Mix (4368708, ThermoFisher Scientific). Primers for HA (5’-AAACTCTTCGCGGTCTTTCCA-3’ sense sequence and 5’-GATAAGGTAGCTTGGGCTGC-3’ antisense sequence), Cre (see primers used for mice genotyping) and ACTB (5’-TACCACAGGCATTGTGATGG-3’ sense sequence and 5’-AATAGTGATGACCTGGCCGT-3’ antisense sequence) were used. Relative HA mRNA expression was determined with 2^-ΔΔCT^ calculation, taking ACTB as reference gene and cDNA from one conserved mouse that received i.v injection of AdCre (AdCre i.v) as positive control. Relative Cre mRNA expression was determined with 2^-ΔCT^ calculation, taking ACTB as reference gene.

### ELISA test

Blood sample clotted for 1h at room temperature followed by centrifugation at 3000g for 10min and serum was harvested and stored at −80°C. For detection of anti-HA antibodies, wells were coated with 1µg/mL of HA protein (11684-V08H, SinoBiological) diluted in coating buffer (Na_2_CO_3_ 0.05M; NaHCO_3_ 0.05M; pH 9.2) and plate was incubated at 4°C overnight. Wells were washed and saturated with dilution buffer (PBS 1X; Tween 20 0.05%; BSA 1%) for 2h at 37°C. Wells were washed, serum samples were diluted by 500, 1000 and 5000 and added to wells in duplicate, and plate was incubated at 37°C for 2h. Wells were washed and 0,4ng/mL of detection antibody (peroxidase-conjugated goat anti-mouse IgG+IgM (H+L) polyclonal antibody, 115-036-068, Jackson ImmunoResearch) was added and plate was incubated at 37°C for 1h. Finally, wells were washed and 50µL of TMB Substrate Reagent (555214, BD Biosciences) was added for revelation step. Reaction was stopped 1min30sec after with H_2_SO_4_ 0.5M. Optical density (OD) at 450nm and 620nm were determined using a Spark® 10M Infinite M200 Pro plate reader (TECAN). Analysis was performed by subtract OD_620nm_ to OD_450nm_ value, calculate the mean of each duplicate and subtract the mean of blank wells to the mean of samples. Then, OD values of each dilution of sample were normalized to OD values of each corresponding dilution of positive control serum sample from one conserved AdCre i.v mouse. Finally, for each sample, mean of normalized OD_450nm_ for each dilution was calculated and analyzed.

### Cell preparation, tetramer enrichment and staining and flow cytometry

Splenocytes were isolated by mechanical dissociation of spleen in red blood cell lysis buffer (NH_4_Cl 155mM; KHCO_3_ 10mM; EDTA 1mM; Tris 17mM per 1L of sterilized water). Liver non-parenchymal cells (NPCs) were isolated as previously described^67^ from liver sample perfused with HBSS 1X. Briefly, livers were digested with collagenase IV, then NPCs were enriched by Percoll density gradient centrifugation and red blood cells lysis. 20.10^6^ splenocytes and 3.10^6^ NPCs were stained with both I-A^d^-HA peptide (HNTNGVTAACSHE) and I-E^d^-HA peptide (SFERFEIFPKE) PE-labelled tetramers (NIH Tetramer Core Facility; Atlanta, USA) at room temperature during 1h. Cells were washed and stained with magnetic anti-PE microbeads (130-048-801, Miltenyi) at 4°C during 15min. Cells were washed and enriched using magnetic MS columns (130-042-201, Miltenyi). Positive fraction was kept for HA-specific CD4^+^ T cell detection. Negative fraction was stained with H-2K^d^-HA peptide (PEYWEEQTQRAKSD) PE-labelled tetramer (NIH Tetramer Core Facility; Atlanta, USA) at room temperature during 1h for HA-specific CD8^+^ T cell detection. All cells were washed and viability staining (LIVE/DEAD^TM^ Fixable Aqua Dead Cell Stain kit, L34957, ThermoFisher Scientific) was performed at 4°C during 15min, protected from light. Cells were washed and extracellular staining were performed at 4°C during 20min protected from light, using fluorochrome-coupled anti-mouse antibodies (Supplementary table 1). Fluorescence was measured on BD FACSCantoII or BD FACSAriaII (BD Biosciences; Mountain View, USA). HA-specific memory CD4 (or CD8) T lymphocytes were defined as: LIVE/DEAD^-^ CD19^-^ CD4^+^ (or CD8^+^) CD44^high^ tetramer^+^ cells. FlowJo software (TreeStar Inc) was used to analyze flow cytometry data.

### 3’-end bulk RNA sequencing

50 up to 200 HA-specific memory CD4^+^ T cells were FACS sorted. RNA extraction, reverse transcription and PCR steps were directly performed after sorting according to the SMART-Seq® v4 Ultra® Low Input RNA Kit protocol (Takara Bio, USA). Amplified complementary DNA were transferred to the Benaroya Research Institute (Seattle, USA) to perform a 3’-end bulk RNA sequencing. To normalize, trimmed-mean of M values (TMM) normalization on the raw gene counts was performed. Additionally, because many genes are not expressed in any of the samples and are therefore uninformative, a filtering step was performed requiring that genes are expressed with at least 1 count per million total reads in at least 10% of the total number of libraries. For the differential gene expression analysis, we used the following threshold: fold change (FC) > 2 and adjusted p value < 0.1.

### Histological analysis

Liver lobes were fixed in formol 4% (VWR) for 24h at room temperature. Dehydration in differential absolute ethanol baths, embedding in paraffin and staining of paraffin-embedded sections (3µm) with hematoxylin-phloxin-saffron (HPS) coloration were performed by the IBISA MicroPICell facility (Biogenouest; Nantes, FRANCE). Slides were scanned and observed using NanoZoomer (Hamamatsu) and NDP Scan software. Histological analysis was performed thanks to a liver inflammation scoring method based on established hepatitis grading^68^, simplified to portal inflammation score (0 to 4), lobular inflammation score (0 to 4) and presence of interface hepatitis (0 to 1), giving a minimum score of 0 and maximum score of 9. Liver inflammation scoring was performed blindly.

### Western blot

Total proteins were extracted from liver samples via RIPA buffer treatment. 25µg of protein were denatured at 95°C for 5min in Laemmli Sample Buffer (161747, Bio-Rad) with DTT 0.1M. Preparation was separated by SDS-PAGE on Mini-PROTEAN TGX Precast Protein Gels (4561036, Bio-Rad) in migration buffer (Tris 15g/L; Glycine 72g/L; SDS 10g/L) and transferred onto a PVDF membrane with the Trans-Blot Turbo Transfer System (Bio-Rad). The membrane was blocked for 2h with a blocking solution (TBS; Tween 20 0.1%; Skim milk 5%), followed by overnight incubation at 4°C with anti-HA antibody (rabbit polyclonal antibody; 11684-T62, Sinobiological) and anti-mouse β-actin antibody (mouse monoclonal antibody; 3700, Cell Signaling) as primary antibodies. The membrane was washed and incubated 1h at room temperature with peroxidase-conjugated donkey anti-rabbit IgG (H+L) antibody (E-AB-1080-120, Cliniscience) and peroxidase-conjugated goat anti-mouse IgG+IgM (H+L) antibody (115-036-068, Jackson ImmunoResearch). Revelation was performed by Electrochemioluminescence Super Signal West Pico (34577, ThermoFisher Scientific) according to the manufacturer instructions. Imaging and analysis of western blots were performed on the ChemiDoc MP Imaging System (Bio-Rad).

### Statistical analysis

Statistical comparisons were performed using GraphPad Prism software V.5 (GraphPad Software, La Jolla, CA, USA) and FaDA^69^ (https://shiny-bird.univ-nantes.fr/app/Fada). P value <0.05 after adjustment were considered significant.

## Acknowledgements

We thank the biological resource centre for biobanking (CHU Nantes, Hôtel Dieu, Centre de ressources biologiques (CRB), Nantes, F-44093, France (BRIF: BB-0033-00040)). We thanks all the member of the HEPATIMGO network. We thank the BIRD Core facility of the SFR Santé F. Bonamy (Université de Nantes, Inserm UMS016, CNRS UMS3556). We thank the Genomics core facility and the Computational Biology, Biostatistics and Modeling (CB2M) hub of CIML. Centre de Calcul Intensif d’Aix-Marseille is acknowledged for granting access to its high performance computing resources. Supported by the “Agence Nationale de la Recherche” (ANR-19-CE17-0024), the patient association “Association pour la lutte contre les maladies inflammatoires du foie et des voies biliaires” (ALBI), the “Fondation Maladies Rares” (including the « High throughput sequencing and rare diseases » program), the “Région pays de la loire”, the LabEx IGO program (n° ANR-11-LABX-0016) funded by the «Investment into the Future» French Government program managed by the “Agence Nationale de la Recherche” (ANR). This work was supported by institutional grants from INSERM, CNRS and Aix-Marseille University to the CIML.

## Author Contributions

A. C. and T. G. performed the experiments, analyzed the data and wrote the manuscript; C. D. analyzed the data; L. G and S. A. performed the experiments, P-J. G. performed the experiments and analyzed the data; M. B. analyzed the data; R. D. analyzed the data; C. S. performed the experiments and analyzed the data; A. D. performed the experiments; P. P-G. supervised data analysis, analyzed the data and wrote the manuscript; C. C. managed human AILD samples collection and patient data base; L. B. and J-P. J. performed experiments; C. F. performed experiments; A. I., M. K., E. B-J., L. E. A. L., C. S., M. S. and F. T. provided human AILD samples and critical insight in AILD pathology; F. V. managed human AILD samples collection; L. B. performed the experiments; S. B. provided critical insight the study design, W. W. K. provided the MHC class II tetramer technology; J-F. M. provided human liver AILD biopsies and critical insight in histology analysis; A. L. provided human AILD samples and critical insight in AILD pathology; J. P. supervised data analysis; M. B. supervised data analysis and critical insight TCR sequencing; J. G. provided human AILD samples, critical insight in AILD pathology and direction in the study design; S. C. designed the study, supervised data analysis and wrote the manuscript; P. M. designed the study, supervised data analysis and wrote the manuscript, A. R. designed the study, supervised data analysis, performed the experiments, analyzed the data and wrote the manuscript. All authors reviewed and approved the manuscript.

## Conflicts of Interest

The authors declare no conflict of interest related to this work.

## Data Availability

Single-cell RNA sequencing raw data from the different experiments analyzed in this article have been deposited in the NCBI GEO repository under accession numbers XXXXX. Processed and annotated single-cell RNA-sequencing datasets analyzed in this article are available in the following repository: XXXXX. Scripts used for analyzing the datasets and produce the figures of this article are available in the following repository: GitHub/XXXX.

---

**Supplementary figure 1:**
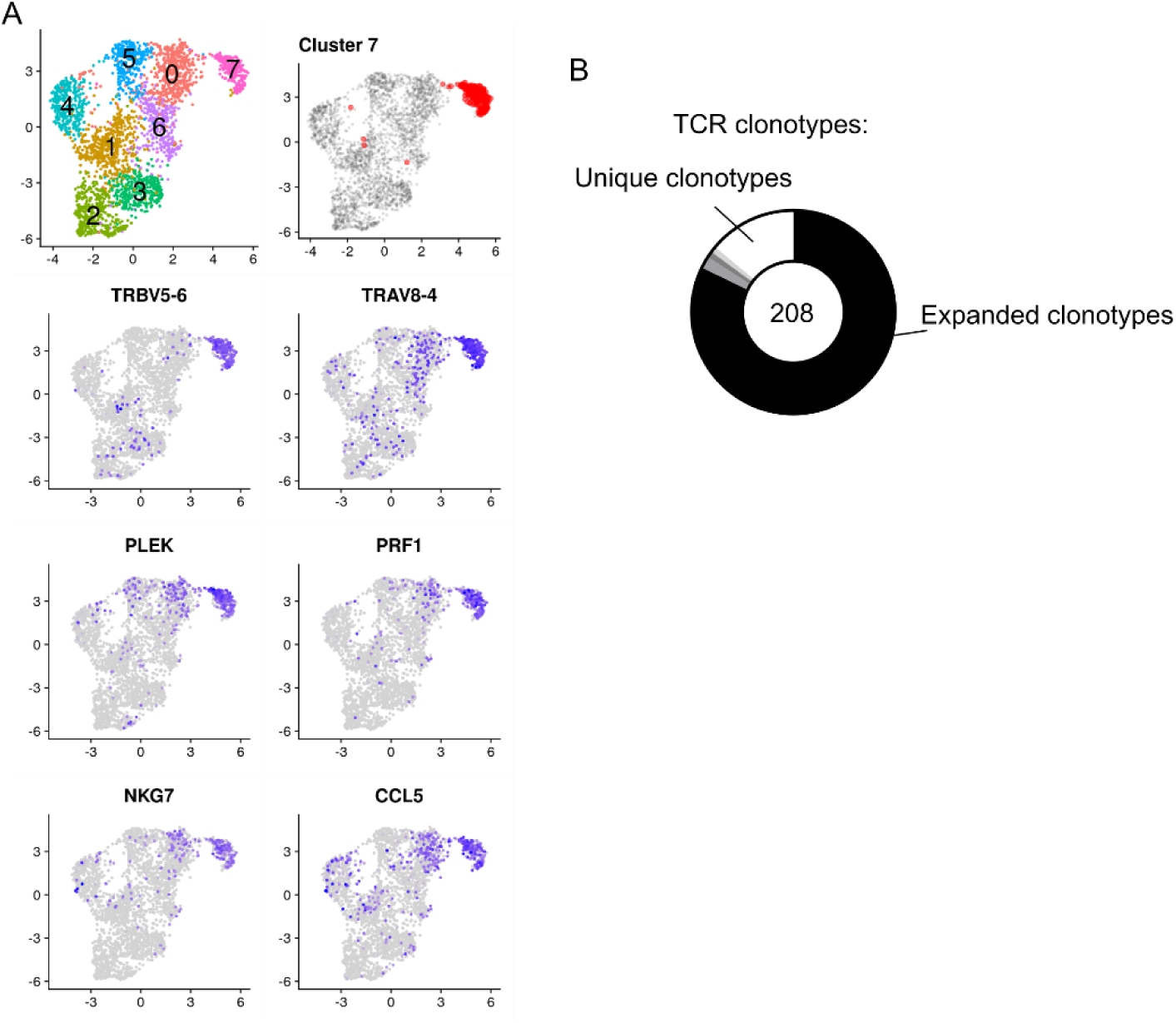
Characterization of the cluster 7 of PDCE2-specific CD4 T cells. (A) UMAP representation of selected gene markers. (B) TCRαβ clonal diversity of antigen-specific single T cells for cluster 7 cells. Numbers indicate the number of single cells analyzed with a TCRαβ sequence. Black and grey sectors indicate the proportion of TCRαβ clones (clonotype common to ≥ 2 cells) within single-cells analyzed; white sector: unique clonotypes.

**Supplementary figure 2:**
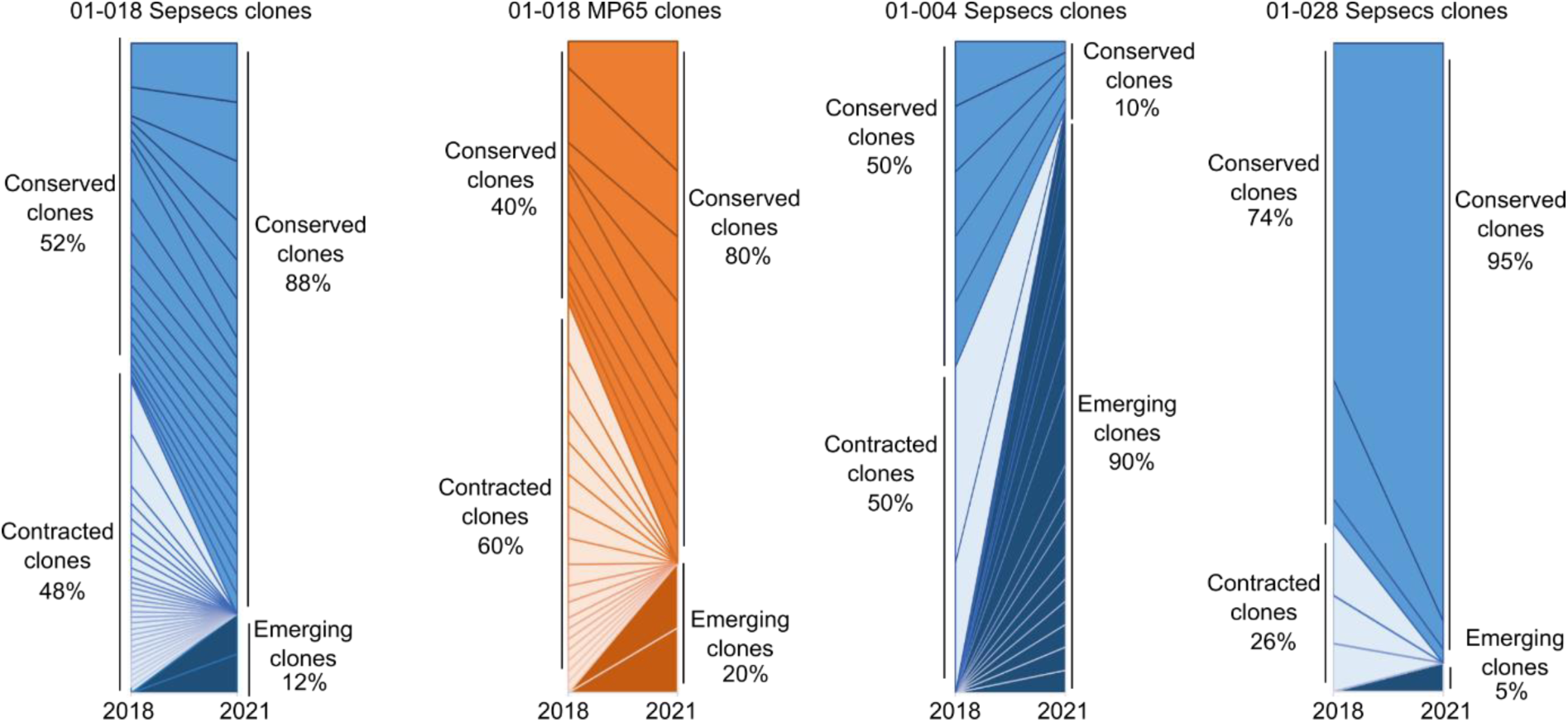
Longitudinal analysis of Sepsecs-specific clonotypes. Representation of the proportion of Sepsecs- and MP65-clonotypes between 2018 and 2021 in the blood of three patients.

**Supplementary figure 3:**
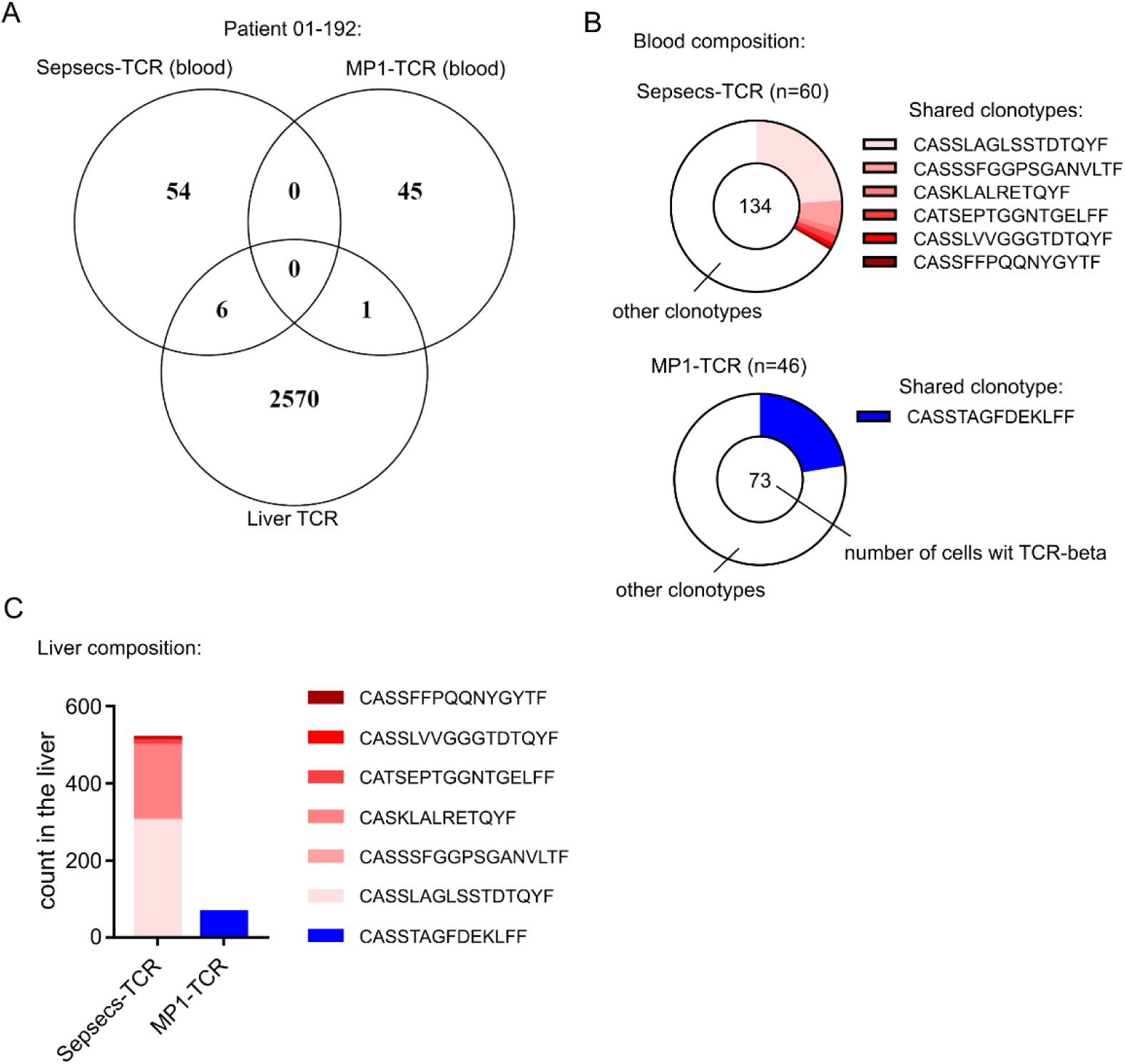
Sepsecs-clonotypes are enriched in the liver of an SLA^+^ patient. (A) Shared TCR between Sepsecs-specific CD4 T cells, MP1-specific CD4 T cells and the liver biopsy. (B) Proportion of shared TCR with the liver (colored sectors) within TCRαβ clonal diversity of Sepsecs- or MP1-specific CD4 T cells. (C) Liver frequency (count) of shared-TCRβ sequences.

**Supplementary figure 4:**
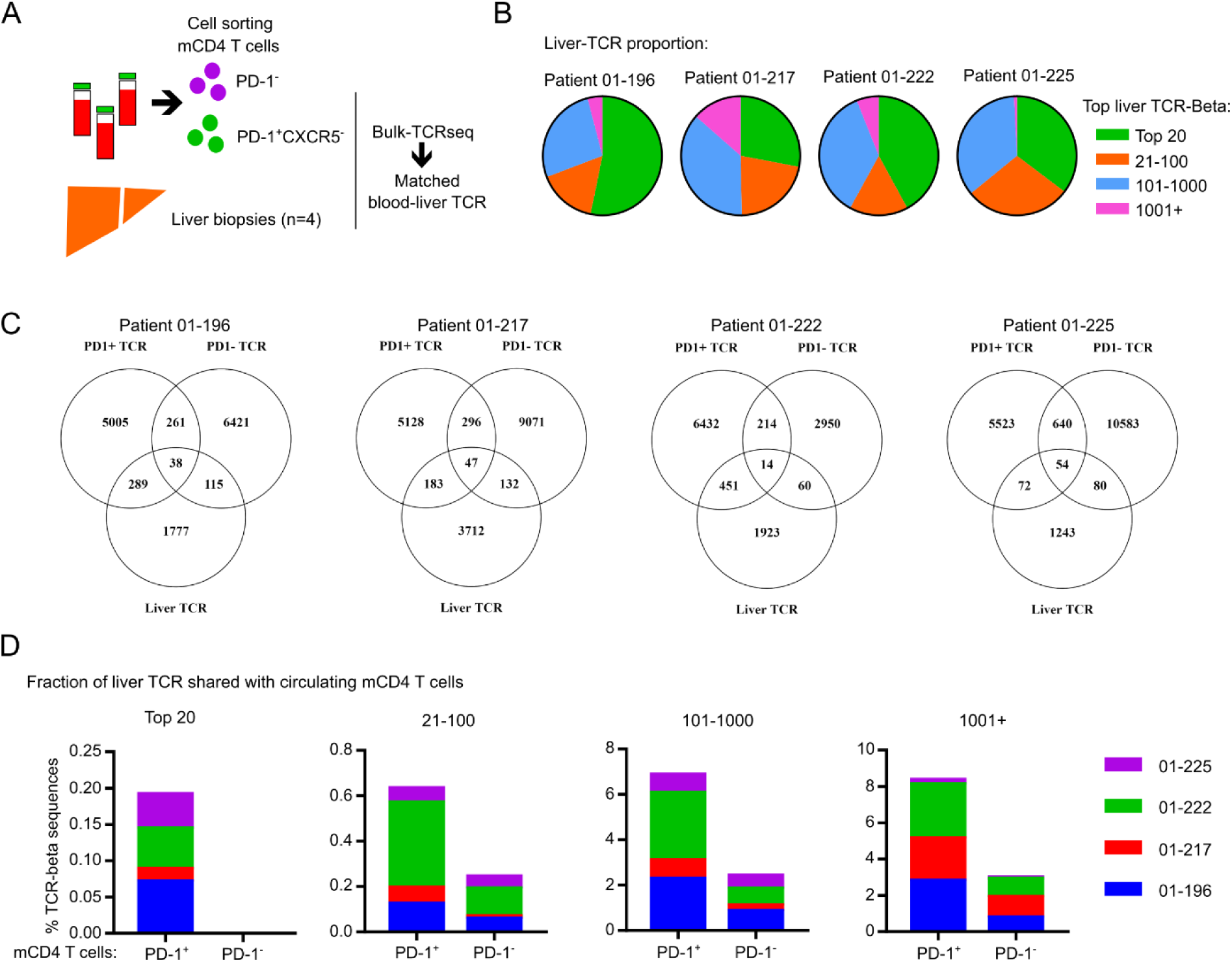
Shared TCRβ sequences between Blood PD-1^+^ or PD-1^-^ CD4 T cells and the liver biopsies from four distinct patients. (A) Experimental design for bulk TCRβ sequencing. (B) Pie chart showing the proportion of the TCRβ repertoire occupied by the top 20, 100, 1000 or 1001+ clones. (C) Shared TCRβ sequences between Blood PD-1^+^ or PD-1^-^ mCD4 T cells and the liver biopsies from the four distinct patients. (D) Analysis of the percentage of TCRβ clonotypes shared between the top 20, 100, 1000 or 1001+ liver clones and the circulating PD-1^+^ CXCR5^-^ (PD-1^+^) or PD-1^-^ mCD4 T cells (PD-1^-^) from four distinct AILD patients.

**Supplementary figure 5:**
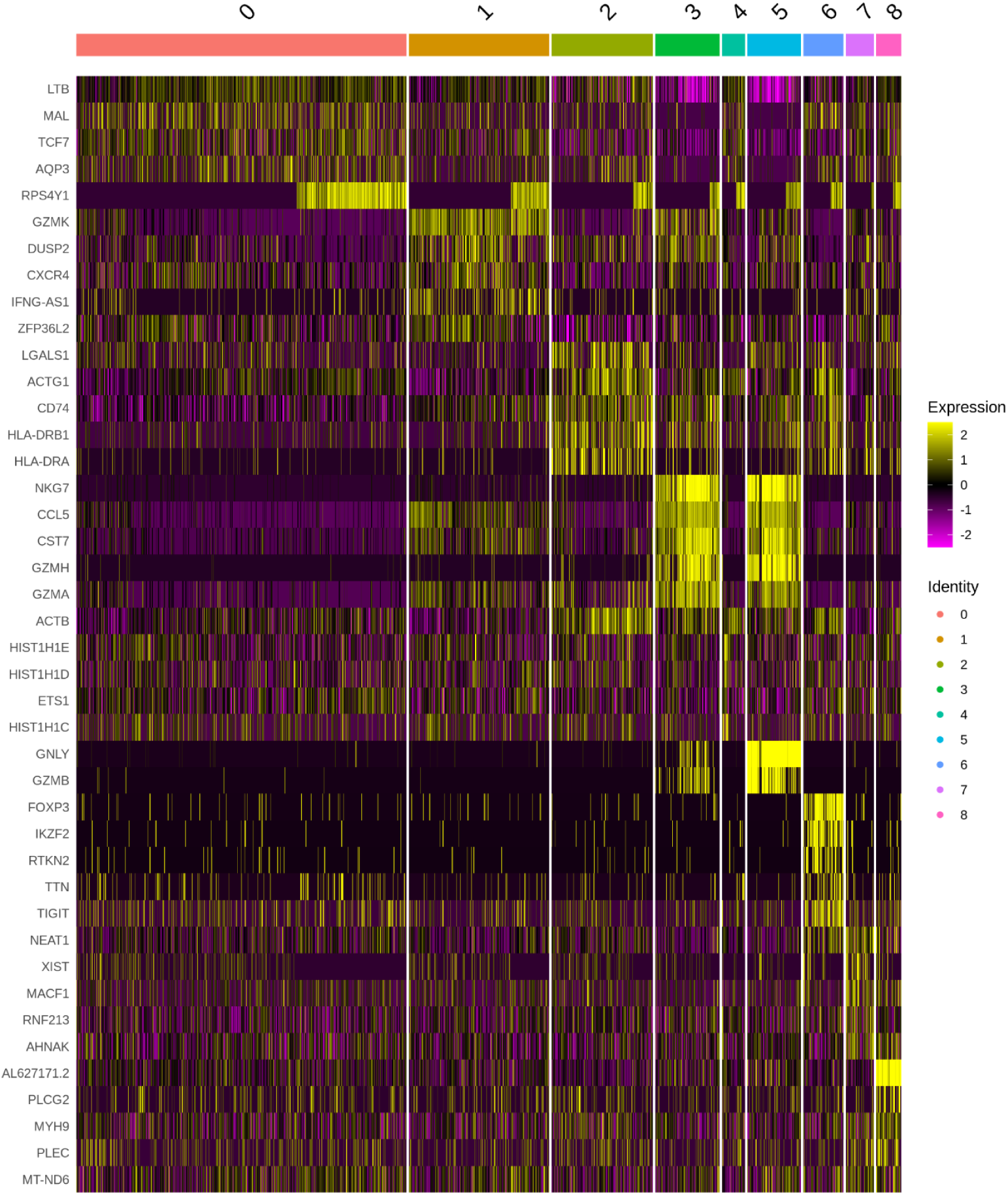
Single cell gene expression heatmap for top 5 marker genes of PD-1^+^ CXCR5^-^ memory CD4 T cells.

**Supplementary figure 6:**
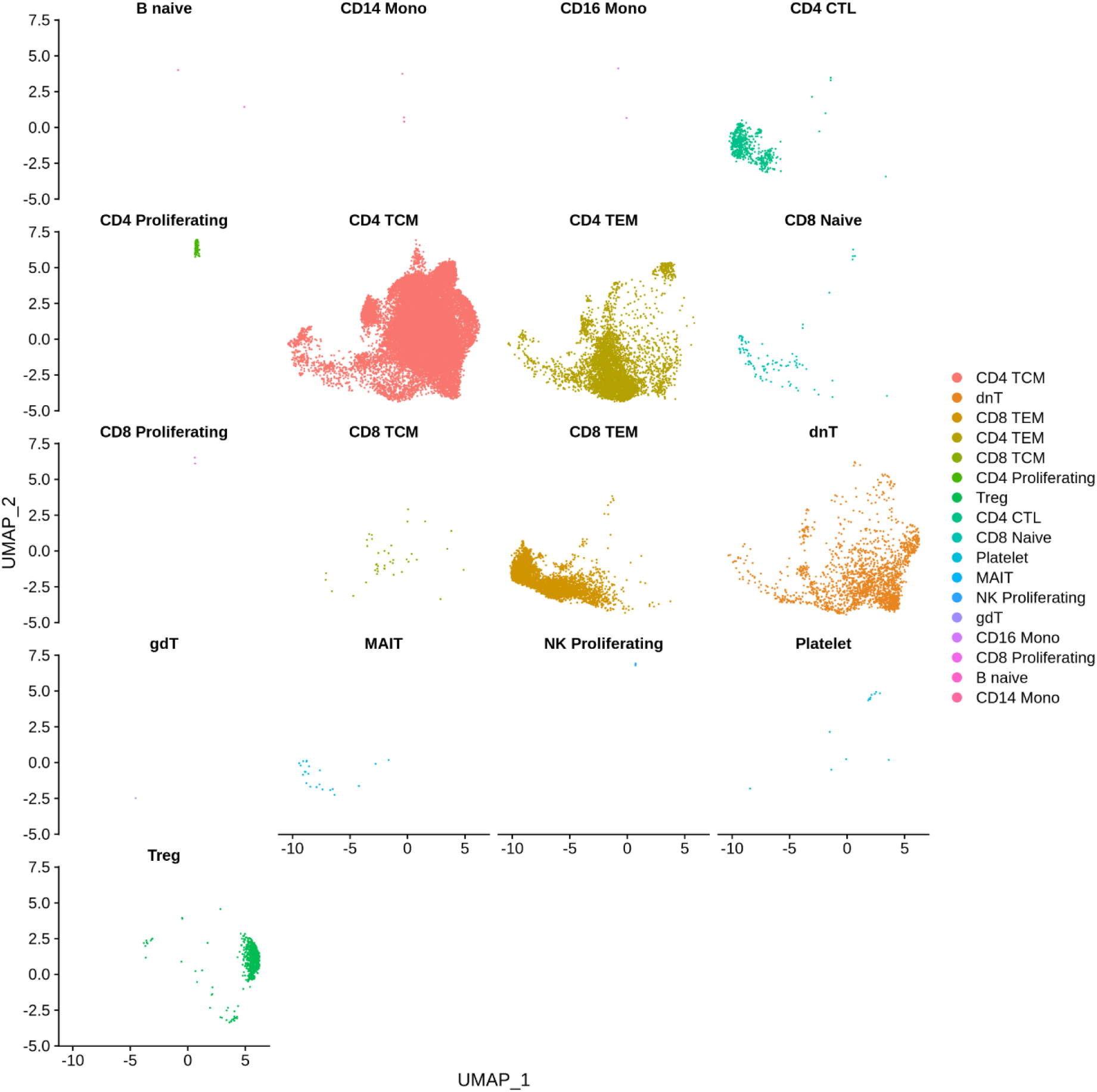
Unsupervised azimuth annotation of scRNA-seq data from PD-1^+^ CXCR5^-^ memory CD4 T cell subsets.

**Supplementary figure 7:**
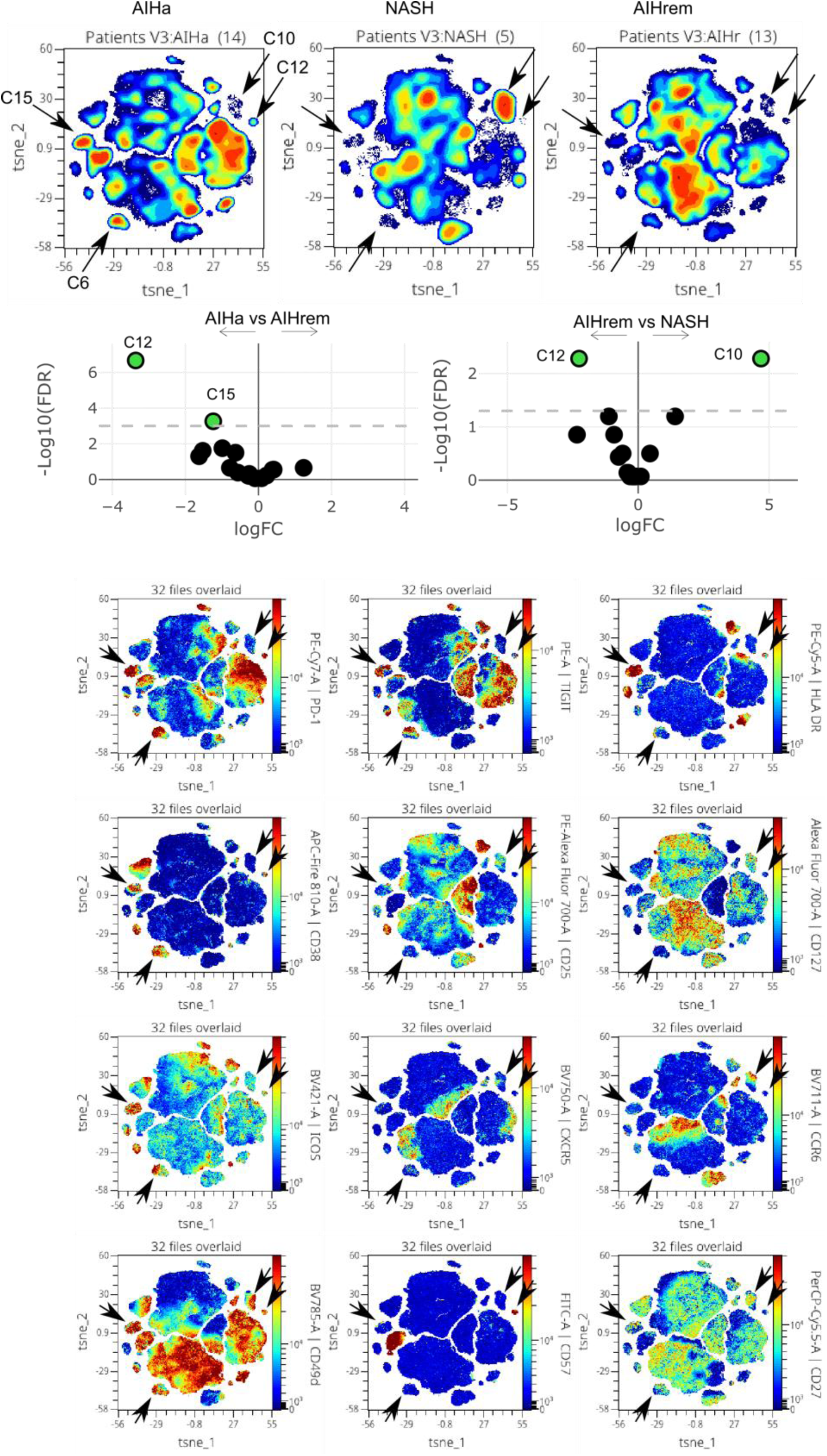
Detailed unsupervised flow cytometry analysis of active AIH (AIHa), NASH and AIH in remission (AIHrem) patients and expression of indicated markers on the opt-SNE representation of blood memory CD4 T cell subsets.

**Supplementary figure 8:**
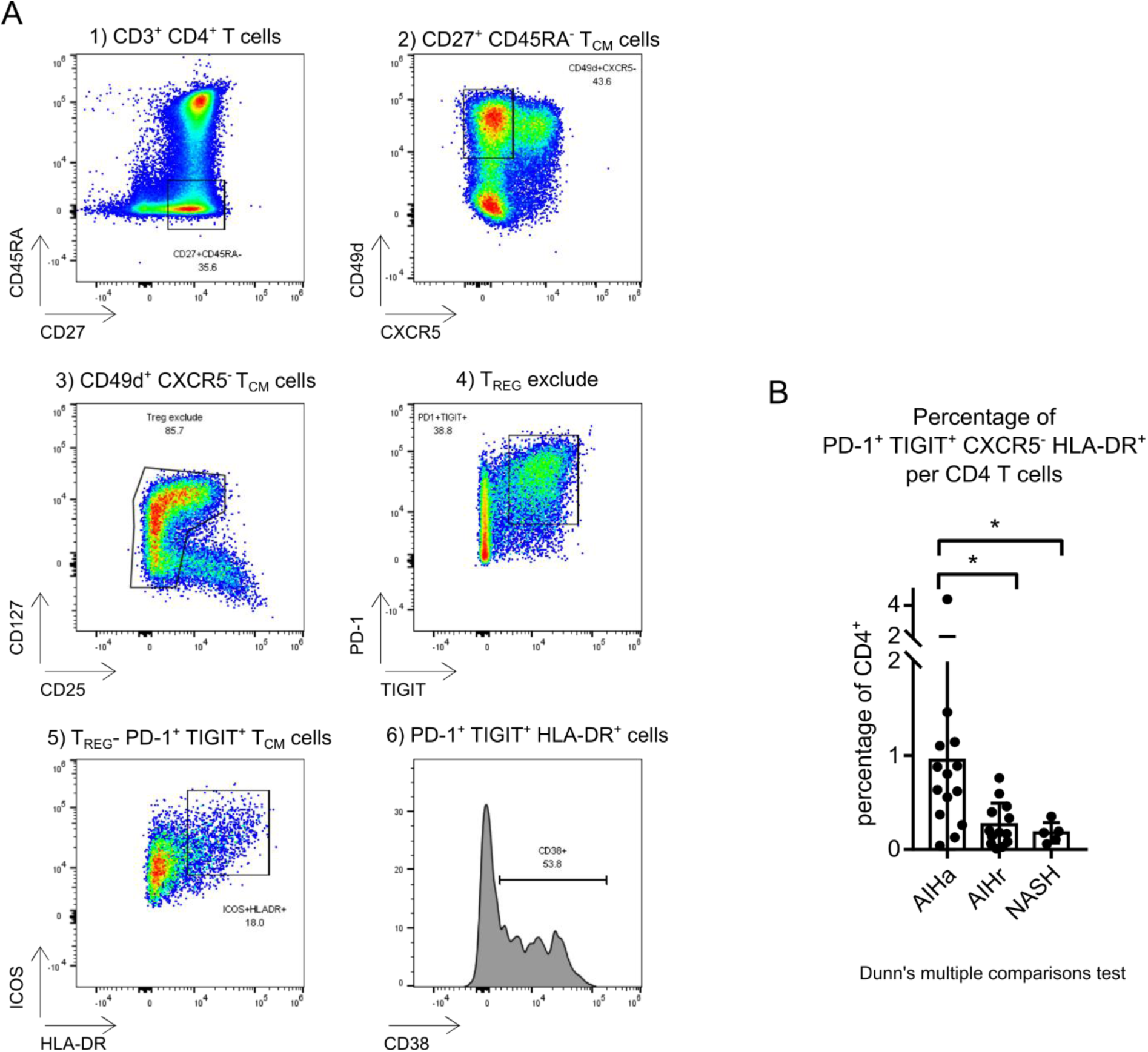
Supervised identification of the cluster 08 (PD-1^+^ CXCR5^-^ TIGIT^+^ HLA-DR^+^). (A) dot plot representation of the gating strategy. (B) Frequency of PD-1^+^ CXCR5^-^ TIGIT^+^ HLA-DR^+^ CD4 T cells per total CD4^+^ T cells in the blood of 5 NASH, 14 active AIH (AIHa), and 13 AIH patients in remission (AIHr) under treatment (<2 years, n=4; >2years, n=12).

**Supplementary figure 9:**
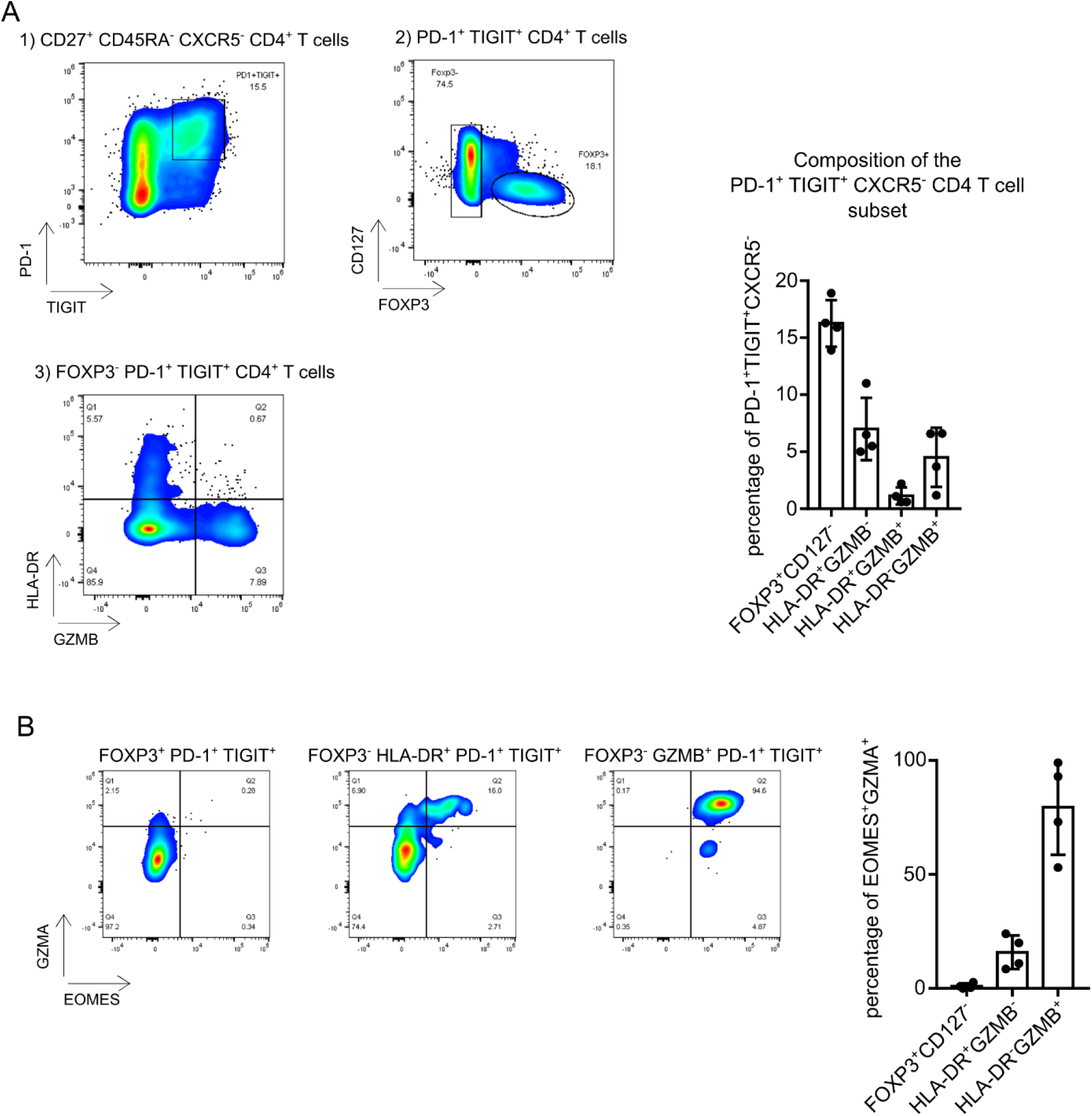
Intracellular characteristics of PD-1^+^ CXCR5^-^ TIGIT^+^ memory CD4 T cells. (A) Foxp3, Granzyme B (GZMB) and HLA-DR expression within PD-1^+^ CXCR5^-^ TIGIT^+^ memory CD4 T cells. (B) Eomes and GZMA expression by PD-1^+^ CXCR5^-^ TIGIT^+^ FOXP3^+^ (FOXP3^+^ CD127^-^); PD-1^+^ CXCR5^-^ TIGIT^+^ FOXP3^-^ HLA-DR^+^ (HLA-DR^+^ GZMB^-^) and PD-1^+^ CXCR5^-^ TIGIT^+^ GZMB+ memory CD4 T cells (HLA-DR^-^ GZMB^+^).

**Supplementary figure 10:**
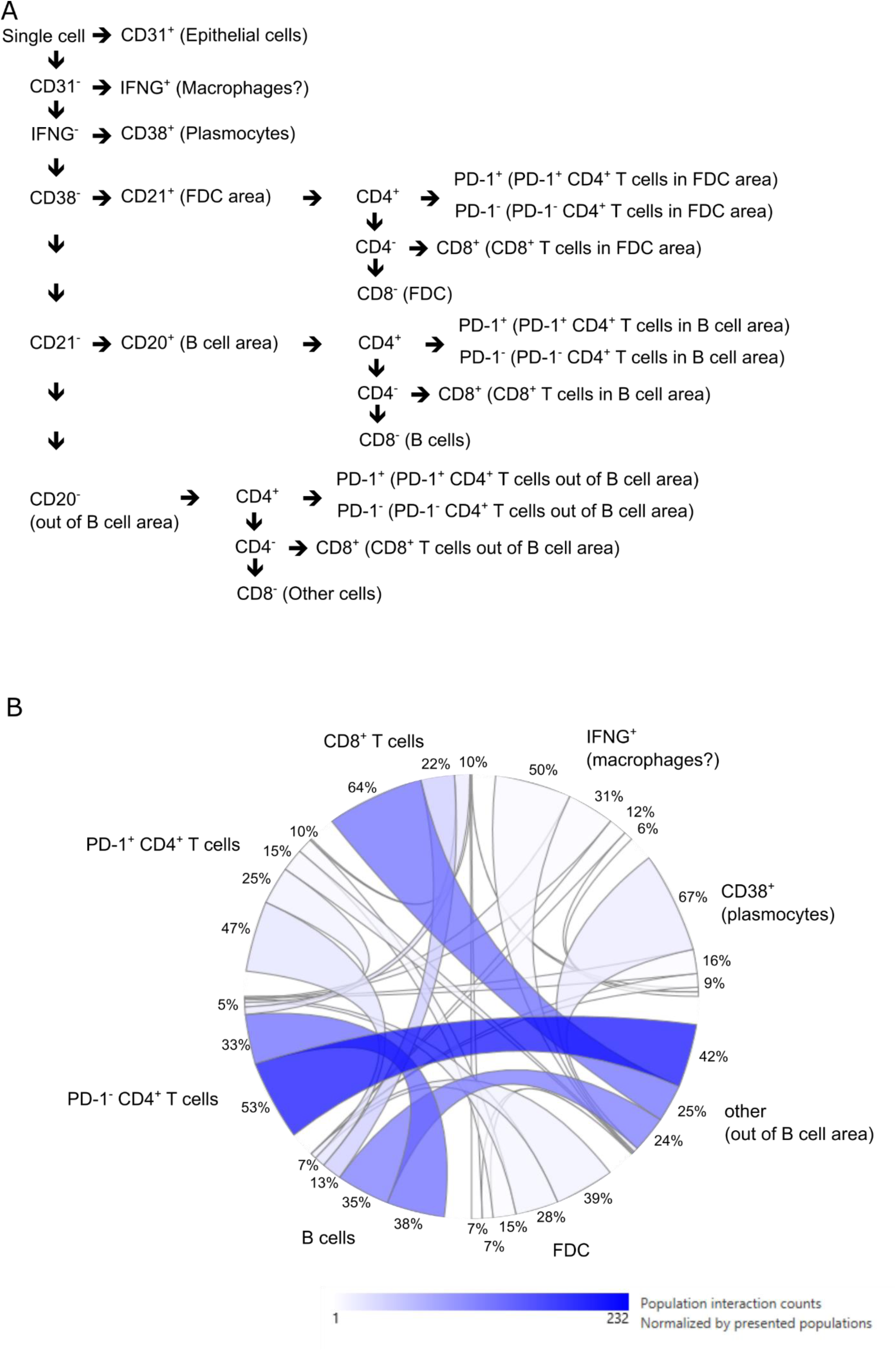
Spatial lineage assignment strategy and proximity analysis. (A) Schematic lineage assignment strategy. (B) Proximity analysis between major cellular subsets identified. Analysis performs with the Multiple Analysis Viewer (MAV) software.

**Supplementary Figure 11:**
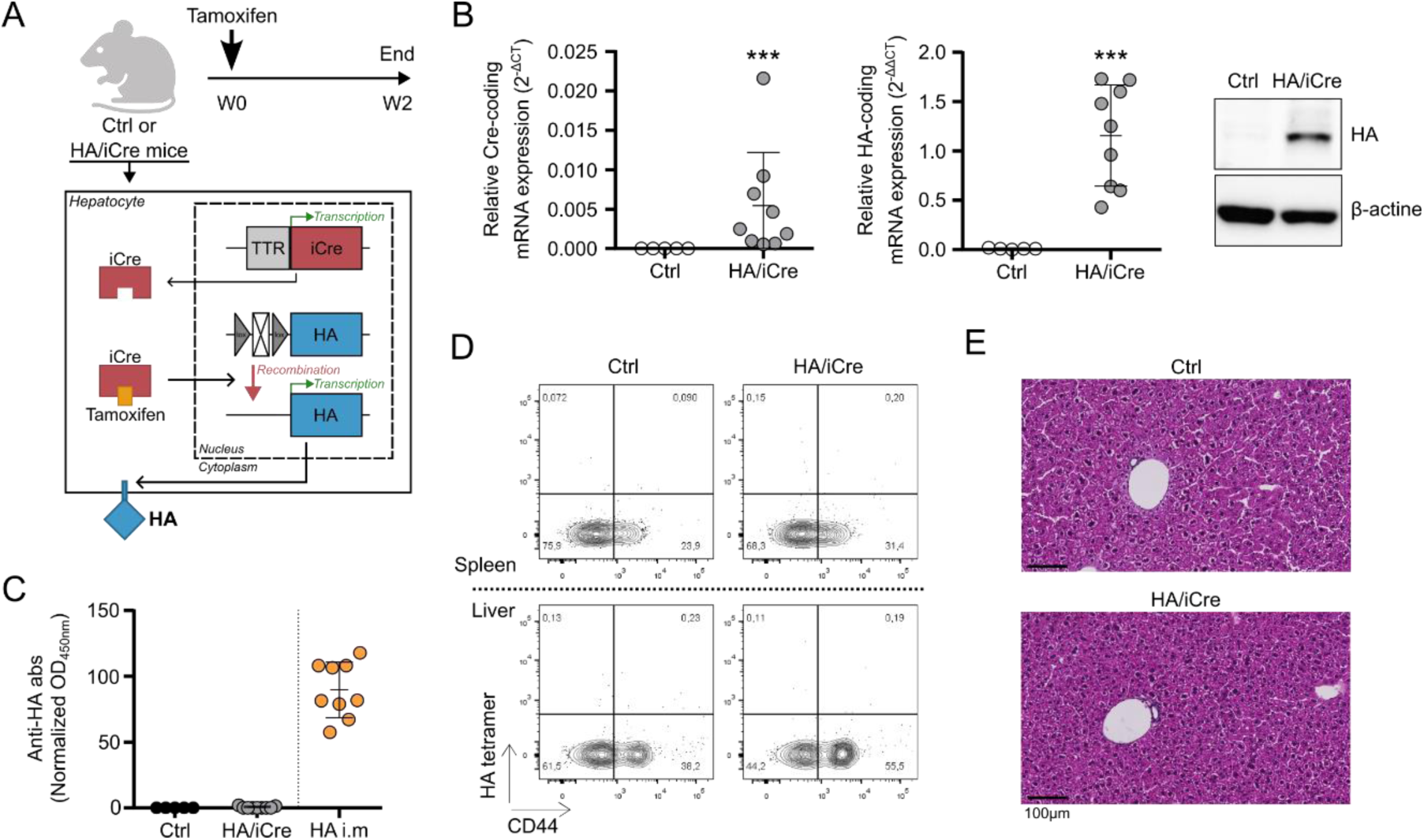
Tamoxifen treatment in non-TCR-transgenic mouse model. (A) Experimental design and schematic illustration for tamoxifen treatment in Rosa26 HA floxed mice (Ctrl) and Rosa26 HA floxed TTR-inducible Cre mice (HA/iCre). (B) Relative Cre mRNA expression (left) and relative HA mRNA expression (middle) in the liver. HA protein detection from total protein extract of liver samples (left). β-actin was used as loading control. (C) Analysis of normalized anti-HA antibody rate in serum of tamoxifen-treated Ctrl and HA/iCre mice. Anti-HA antibody rate of immunized mice (HA i.m) was indicated as positive control. (D) Contour plot representation of HA tetramer staining and CD44 expression in CD4 T cells from spleen (top) and liver (bottom) of tamoxifen-treated Ctrl and HA/iCre mice. (E) Representative pictures of paraffin-embedded liver sections stained with HPS coloration. Black line is used as scale. Mann-Whitney test was used for B. ***: p<0.001.

**Supplementary Table1.**
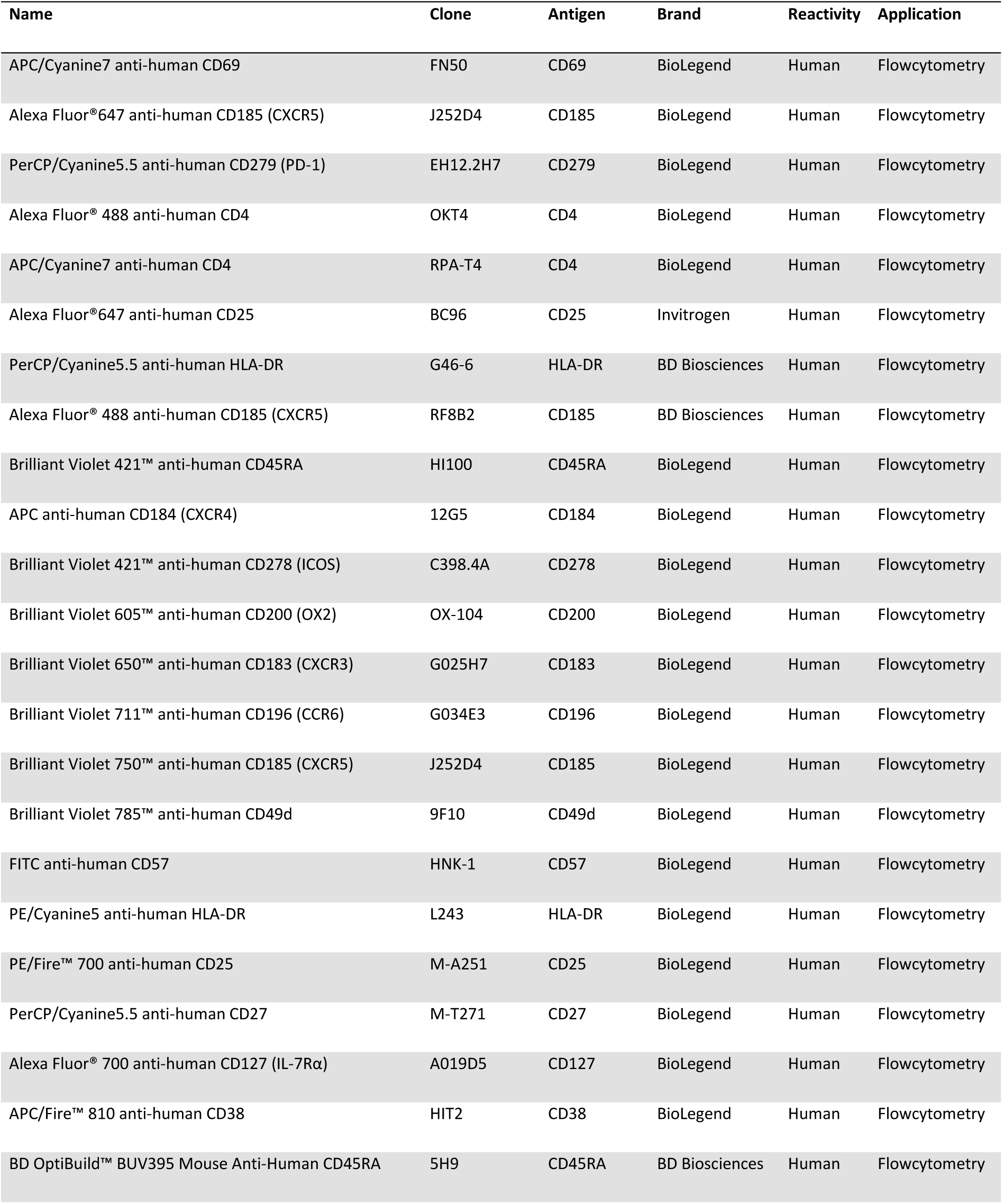
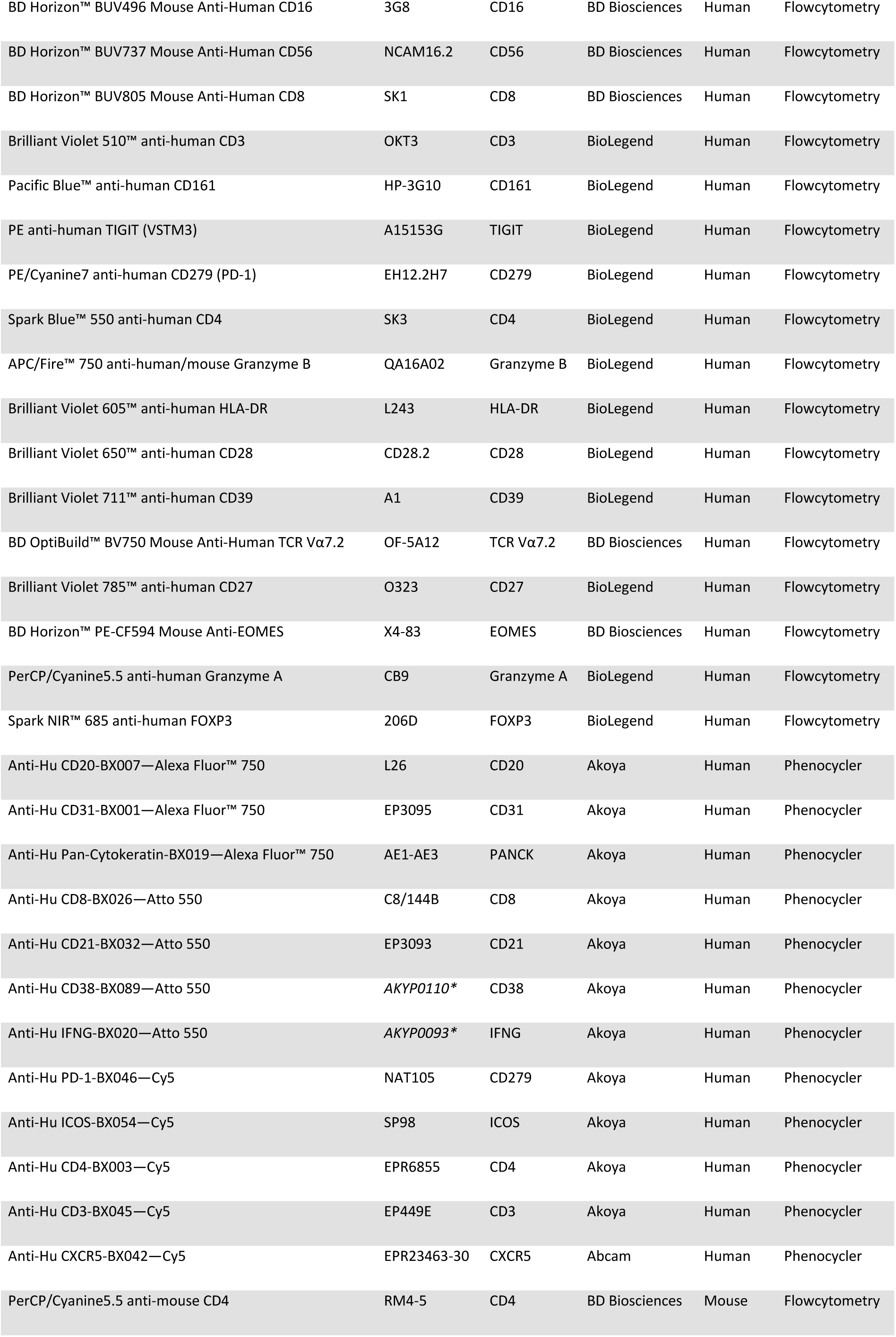
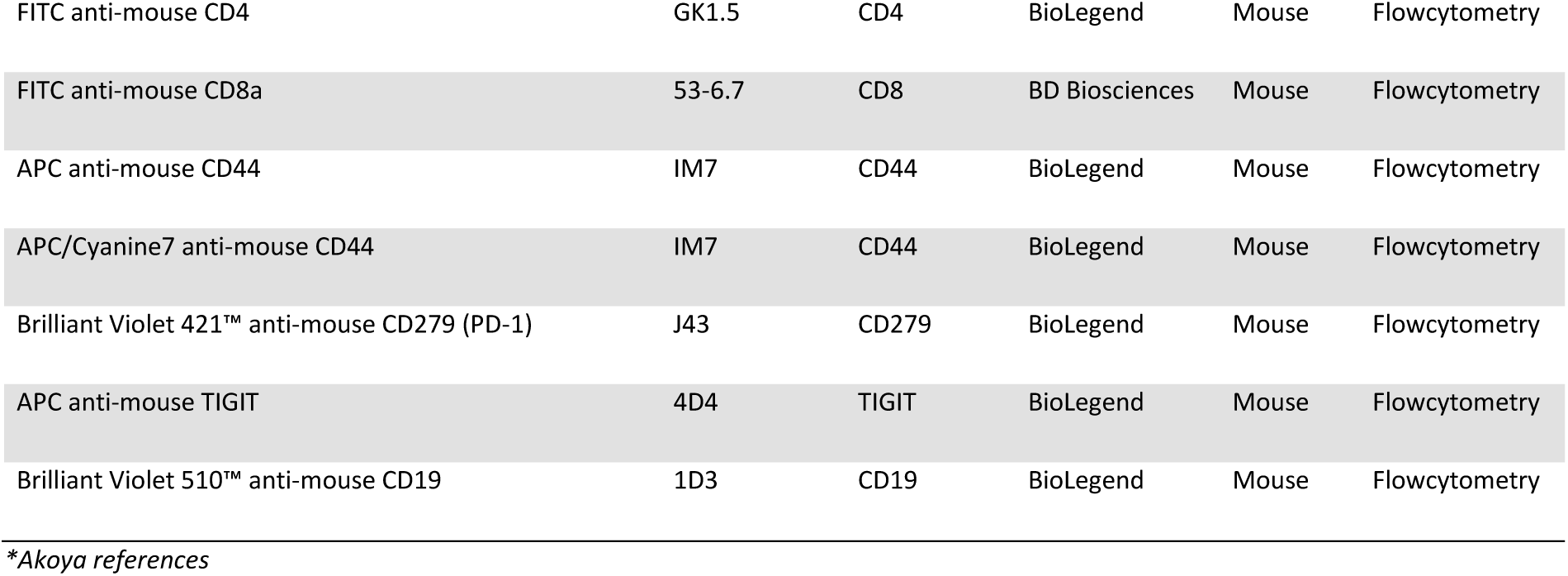
Antibody listing.

| Name | Clone | Antigen | Brand | Reactivity | Application |
| --- | --- | --- | --- | --- | --- |
| APC/Cyanine7 anti-human CD69 | FN50 | CD69 | BioLegend | Human | Flowcytometry |
| Alexa Fluor® 647 anti-human CD185 (CXCR5) | J252D4 | CD185 | BioLegend | Human | Flowcytometry |
| PerCP/Cyanine5.5 anti-human CD279 (PD-1) | EH12.2H7 | CD279 | BioLegend | Human | Flowcytometry |
| Alexa Fluor® 488 anti-human CD4 | OKT4 | CD4 | BioLegend | Human | Flowcytometry |
| APC/Cyanine7 anti-human CD4 | RPA-T4 | CD4 | BioLegend | Human | Flowcytometry |
| Alexa Fluor® 647 anti-human CD25 | BC96 | CD25 | Invitrogen | Human | Flowcytometry |
| PerCP/Cyanine5.5 anti-human HLA-DR | G46-6 | HLA-DR | BD Biosciences | Human | Flowcytometry |
| Alexa Fluor® 488 anti-human CD185 (CXCR5) | RF8B2 | CD185 | BD Biosciences | Human | Flowcytometry |
| Brilliant Violet 421™ anti-human CD45RA | HI100 | CD45RA | BioLegend | Human | Flowcytometry |
| APC anti-human CD184 (CXCR4) | 12G5 | CD184 | BioLegend | Human | Flowcytometry |
| Brilliant Violet 421™ anti-human CD278 (ICOS) | C398.4A | CD278 | BioLegend | Human | Flowcytometry |
| Brilliant Violet 605™ anti-human CD200 (OX2) | OX-104 | CD200 | BioLegend | Human | Flowcytometry |
| Brilliant Violet 650™ anti-human CD183 (CXCR3) | G025H7 | CD183 | BioLegend | Human | Flowcytometry |
| Brilliant Violet 711™ anti-human CD196 (CCR6) | G034E3 | CD196 | BioLegend | Human | Flowcytometry |
| Brilliant Violet 750™ anti-human CD185 (CXCR5) | J252D4 | CD185 | BioLegend | Human | Flowcytometry |
| Brilliant Violet 785™ anti-human CD49d | 9F10 | CD49d | BioLegend | Human | Flowcytometry |
| FITC anti-human CD57 | HNK-1 | CD57 | BioLegend | Human | Flowcytometry |
| PE/Cyanine5 anti-human HLA-DR | L243 | HLA-DR | BioLegend | Human | Flowcytometry |
| PE/Fire™ 700 anti-human CD25 | M-A251 | CD25 | BioLegend | Human | Flowcytometry |
| PerCP/Cyanine5.5 anti-human CD27 | M-T271 | CD27 | BioLegend | Human | Flowcytometry |
| Alexa Fluor® 700 anti-human CD127 (IL-7Rα) | A019D5 | CD127 | BioLegend | Human | Flowcytometry |
| APC/Fire™ 810 anti-human CD38 | HIT2 | CD38 | BioLegend | Human | Flowcytometry |
| BD OptiBuild™ BUV395 Mouse Anti-Human CD45RA | 5H9 | CD45RA | BD Biosciences | Human | Flowcytometry |
| BD Horizon™ BUV496 Mouse Anti-Human CD16 | 3G8 | CD16 | BD Biosciences | Human | Flowcytometry |
| BD Horizon™ BUV737 Mouse Anti-Human CD56 | NCAM16.2 | CD56 | BD Biosciences | Human | Flowcytometry |
| BD Horizon™ BUV805 Mouse Anti-Human CD8 | SK1 | CD8 | BD Biosciences | Human | Flowcytometry |
| Brilliant Violet 510™ anti-human CD3 | OKT3 | CD3 | BioLegend | Human | Flowcytometry |
| Pacific Blue™ anti-human CD161 | HP-3G10 | CD161 | BioLegend | Human | Flowcytometry |
| PE anti-human TIGIT (VSTM3) | A15153G | TIGIT | BioLegend | Human | Flowcytometry |
| PE/Cyanine7 anti-human CD279 (PD-1) | EH12.2H7 | CD279 | BioLegend | Human | Flowcytometry |
| Spark Blue™ 550 anti-human CD4 | SK3 | CD4 | BioLegend | Human | Flowcytometry |
| APC/Fire™ 750 anti-human/mouse Granzyme B | QA16A02 | Granzyme B | BioLegend | Human | Flowcytometry |
| Brilliant Violet 605™ anti-human HLA-DR | L243 | HLA-DR | BioLegend | Human | Flowcytometry |
| Brilliant Violet 650™ anti-human CD28 | CD28.2 | CD28 | BioLegend | Human | Flowcytometry |
| Brilliant Violet 711™ anti-human CD39 | A1 | CD39 | BioLegend | Human | Flowcytometry |
| BD OptiBuild™ BV750 Mouse Anti-Human TCR Vα7.2 | OF-5A12 | TCR Vα7.2 | BD Biosciences | Human | Flowcytometry |
| Brilliant Violet 785™ anti-human CD27 | O323 | CD27 | BioLegend | Human | Flowcytometry |
| BD Horizon™ PE-CF594 Mouse Anti-EOMES | X4-83 | EOMES | BD Biosciences | Human | Flowcytometry |
| PerCP/Cyanine5.5 anti-human Granzyme A | CB9 | Granzyme A | BioLegend | Human | Flowcytometry |
| Spark NIR™ 685 anti-human FOXP3 | 206D | FOXP3 | BioLegend | Human | Flowcytometry |
| Anti-Hu CD20-BX007—Alexa Fluor™ 750 | L26 | CD20 | Akoya | Human | Phenocycler |
| Anti-Hu CD31-BX001—Alexa Fluor™ 750 | EP3095 | CD31 | Akoya | Human | Phenocycler |
| Anti-Hu Pan-Cytokeratin-BX019—Alexa Fluor™ 750 | AE1-AE3 | PANCK | Akoya | Human | Phenocycler |
| Anti-Hu CD8-BX026—Atto 550 | C8/144B | CD8 | Akoya | Human | Phenocycler |
| Anti-Hu CD21-BX032—Atto 550 | EP3093 | CD21 | Akoya | Human | Phenocycler |
| Anti-Hu CD38-BX089—Atto 550 | AKYP0110* | CD38 | Akoya | Human | Phenocycler |
| Anti-Hu IFNG-BX020—Atto 550 | AKYP0093* | IFNG | Akoya | Human | Phenocycler |
| Anti-Hu PD-1-BX046—Cy5 | NAT105 | CD279 | Akoya | Human | Phenocycler |
| Anti-Hu ICOS-BX054—Cy5 | SP98 | ICOS | Akoya | Human | Phenocycler |
| Anti-Hu CD4-BX003—Cy5 | EPR6855 | CD4 | Akoya | Human | Phenocycler |
| Anti-Hu CD3-BX045—Cy5 | EP449E | CD3 | Akoya | Human | Phenocycler |
| Anti-Hu CXCR5-BX042—Cy5 | EPR23463-30 | CXCR5 | Abcam | Human | Phenocycler |
| PerCP/Cyanine5.5 anti-mouse CD4 | RM4-5 | CD4 | BD Biosciences | Mouse | Flowcytometry |
| FITC anti-mouse CD4 | GK1.5 | CD4 | BioLegend | Mouse | Flowcytometry |
| FITC anti-mouse CD8a | 53-6.7 | CD8 | BD Biosciences | Mouse | Flowcytometry |
| APC anti-mouse CD44 | IM7 | CD44 | BioLegend | Mouse | Flowcytometry |
| APC/Cyanine7 anti-mouse CD44 | IM7 | CD44 | BioLegend | Mouse | Flowcytometry |
| Brilliant Violet 421™ anti-mouse CD279 (PD-1) | J43 | CD279 | BioLegend | Mouse | Flowcytometry |
| APC anti-mouse TIGIT | 4D4 | TIGIT | BioLegend | Mouse | Flowcytometry |
| Brilliant Violet 510™ anti-mouse CD19 | 1D3 | CD19 | BioLegend | Mouse | Flowcytometry |

*\*Akoya references*

**Supplementary Table 2.** Clinical and biological characteristics of patients with AIH and NASH expressed as mean [95% confidence interval].

| Group | Active AIH (AIHa) | Remission AIH (AIHr) | NASH |
| --- | --- | --- | --- |
|  | n=14 | n=13 | n=5 |
| Age (year) | 50,3 [39,7-61] | 52,5 [39,1-65,8] | 59,2 [33,3-85,1] |
| Female, n (%) | 9 (64%) | 9 (69%) | 3 (60%) |
| IgG (g/L) | 19,5 [15,2-23,8] | 11,4 [9,9-12,9] | 11,9 [6,8-17,1] |
| AST (IU/L) | 307 [121,7-492,1] | 22,6 [17,4-27,9] | 45,9 [23,4-68,4] |
| ALT (IU/L) | 505 [185,6-824,7] | 20,8 [14,4-27,1] | 73,5 [6,2-140,9] |
| Immunosuppressive agents n (%) |  |  |  |
| No, n (%) | 9 (64%) | 0 | 5 (100%) |
| Naïve | 9 | 0 | NA |
| Yes, n (%) | 5 (36%) | 13 (100%) | 0 |
| Azathioprin | 4 | 11 | NA |
| MMF | 1 | 1 | NA |
| Steroids only | 0 | 1 | NA |
| Steroids associated to other therapy | 2 | 3 | NA |

AIH: Autoimmune Hepatitis, NASH: Non Alcoholic Steatohepatitis, AST: Aspartate Aminotransferase, ALT: Alanine
Aminotransferase, NA: non-applicable

## Notes

### Competing Interest Statement

The authors have declared no competing interest.

### Summary of Updates

This version of the manuscript is an updated pre-submission version; the discussion has been reduced and the list of authors corrected.

